# Acetyl-CoA mediated autoacetylation of fatty acid synthase in *de novo* lipogenesis

**DOI:** 10.1101/2022.01.06.475252

**Authors:** Ting Miao, Jinoh Kim, Ping Kang, Hua Bai

## Abstract

*De novo lipogenesis* (DNL) is a highly regulated metabolic process, which is known to be activated through transcriptional regulation of lipogenic genes, including fatty acid synthase (FASN). Unexpectedly, we find that the expression of FASN protein remains unchanged during *Drosophila* larval development when lipogenesis is hyperactive. Instead, acetylation modification of FASN is highly upregulated in fast-growing larvae. We further show that lysine K813 is highly acetylated in developing larvae, and its acetylation is required for upregulated FASN activity, body fat accumulation, and normal development. Intriguingly, K813 is rapidly autoacetylated by acetyl-CoA in a dosage-dependent manner, independent of known acetyltransferases. Furthermore, the autoacetylation of K813 is mediated by a conserved P-loop-like motif (N-xx-G-x-A). In summary, this work uncovers a novel role of acetyl-CoA-mediated autoacetylation of FASN in developmental lipogenesis and reveals a self-regulatory system that controls metabolic homeostasis by linking acetyl-CoA, lysine acetylation, and DNL.

**Graphical Abstract:** 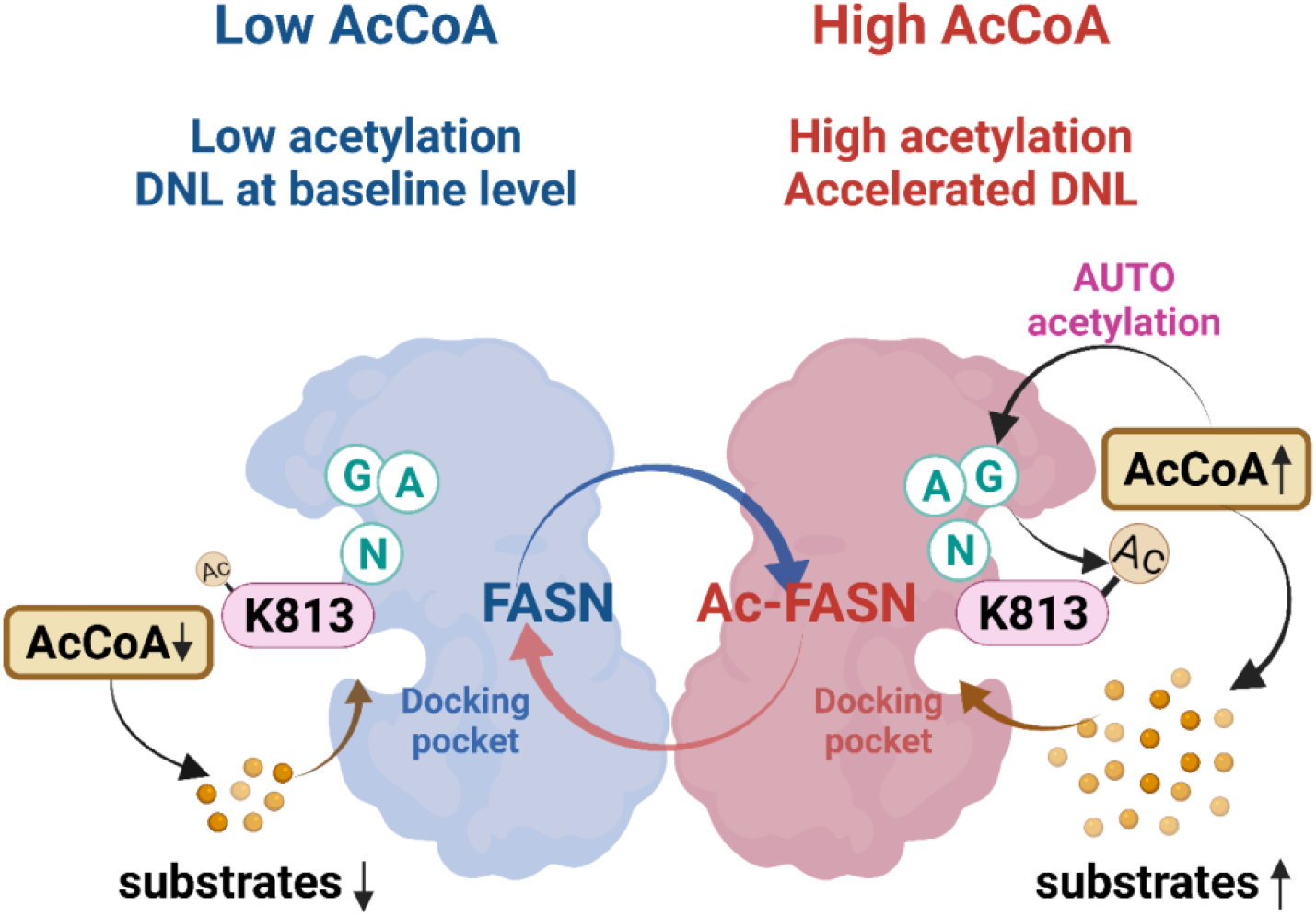

**Highlights:**

- Acetylation modification of FASN, but not protein expression, positively correlates with *de novo* lipogenesis during *Drosophila* larval development
- Site-specific acetylation at K813 residue enhances FASN enzymatic activity
- K813 residue is autoacetylated by acetyl-CoA, independent of KATs
- A novel N-xx-G-x-A motif is required for autoacetylation of K813

## INTRODUCTION

The ability to sense and respond to nutrient availability is essential for organisms to maintain metabolic homeostasis. Dysregulated fatty acid homeostasis is observed in various human diseases, including obesity, diabetes, and most cancers. *De novo lipogenesis* (DNL) is a complex yet highly regulated metabolic process, which converts excess carbohydrates into fatty acids that are then esterified to storage triglyceride (Ameer et al., 2014; Strable and Ntambi, 2010). DNL is often dysregulated in metabolic anomalies. Abnormal upregulation of DNL is an essential contributor to increased fat mass in the pathogenesis of the non-alcoholic fatty liver disease (NAFLD) and diabetes (Chakravarthy et al., 2009; Lin et al., 2005; Loftus, 2000; Nagai et al., 2009; Savage, 2006). DNL is known to be transcriptionally regulated via sterol regulatory element-binding protein 1 (SREBP1c) and carbohydrate-responsive element-binding protein (ChREBP) in response to metabolic and hormonal cues (Horton et al., 2003; Liang et al., 2002; Xu et al., 2013). However, it has been recently proposed that allosteric regulation and post-translational modifications (PTMs) are sometimes used to control metabolic flux, since the timescale of the gene expression is too longer to balance quick turnover of metabolites (Shamir et al., 2016).

Lysine acetylation has recently risen as a novel player that links key metabolites (e.g, acetyl coenzyme A (acetyl-CoA) and nicotinamide adenine dinucleotide (NAD^+^)), cell signaling, and gene regulation (Ali et al., 2018; Menzies et al., 2016). Previous acetylome studies found that almost all metabolic enzymes are acetylated (Nakayasu et al., 2017; Wang et al., 2010a; Zhao et al., 2010), including fatty acid synthase (FASN) (Hennigar et al., 1998; Jin et al., 2010; Lin et al., 2016). FASN, an essential cytosolic enzyme in DNL pathway, catalyzes the biosynthesis of saturated fatty acids from acetyl-CoA, malonyl coenzyme A (malonyl-CoA), and nicotinamide adenine dinucleotide phosphate (NADPH) (Smith, 1994). Recently, FASN has emerged as a novel therapeutic target for the treatment of obesity, diabetes, fatty liver diseases, and cancers (Angeles and Hudkins, 2016; Kridel et al., 2007; Loftus, 2000; Wu et al., 2011). Although FASN is known to be regulated through SREBP1-mediated transcriptional activation (Horton et al., 2003; Liang et al., 2002; Xu et al., 2013), several conflicting results show that there is little correlation between FASN expression and its enzymatic activity (Hennigar et al., 1998; Jin et al., 2010; Najjar et al., 2005; Qureshi et al., 1975; Sabbisetti et al., 2009). These findings suggest a possible involvement of PTMs in the regulation of FASN activities. Phosphorylation and acetylation have been proposed as alternative mechanisms of FASN regulation (Hennigar et al., 1998; Jin et al., 2010; Lin et al., 2016). However, how post-translational modifications (e.g., acetylation) regulate FASN activity and lipogenesis remains largely unknown.

Although protein acetylation is catalyzed mainly by lysine acetyltransferases (KATs), it has been recently reported that acetylation also arises from a nonenzymatic reaction with acetyl-CoA in eukaryotes (Hansen et al., 2019; Olia et al., 2015; Wagner and Payne, 2013). Acetyl-CoA is the acetyl donor for protein acetylation and is a reactive metabolic intermediate that is involved in a variety of metabolic pathways, including DNL. Therefore, acetyl-CoA has emerged as a key signal metabolite linking metabolism to cellular signaling (Pietrocola et al., 2015). The intracellular levels of acetyl-CoA fluctuate in response to both intracellular and extracellular cues (e.g., growth signals and nutrient conditions), which consequently impacts chromatin modifications and transcriptional reprogramming (Pietrocola et al., 2015; Shi and Tu, 2015). Previously, it was thought that nonenzymatic acetylation mainly occurs to mitochondria proteins, as high concentrations of acetyl-CoA and alkaline environment inside mitochondrial matrix favor lysine nucleophilic attack on the carbonyl carbon of acetyl-CoA (Wagner and Payne, 2013). In recent years, the capability of enzyme-independent acetylation of cytosolic proteins was also determined (Kulkarni et al., 2017; Olia et al., 2015). Yet, the mechanism of nonenzymatic acetylation, especially cytosolic proteins, remains elusive.

Here, we show that the expression of *Drosophila* FASN (dFASN) protein remains unchanged during larval development, the stages when lipogenesis is hyperactive. In contrast, acetylation of dFASN at K813 residue is significantly induced in response to increased cellular acetyl-CoA levels, which promotes dFASN enzymatic activity and lipogenesis in fast-growing larvae. Strikingly, we find that acetylation modification of K813 is controlled through a unique KAT-independent mechanism, which involves a novel motif ‘N-xx-G-x-A’. Thus, our findings uncover a novel acetyl-CoA-mediated self-regulatory module that regulates developmental DNL via autoacetylation of dFASN.

## RESULTS

### Acetylation modification of dFASN, but not protein expression, is positively correlated with *de novo* lipogenesis during *Drosophila* larval development

The fruit fly *Drosophila* has been established as an excellent model to study human diseases and metabolic pathways, given the genetic tractability, sophisticated genetic tools, and evolutionarily conserved metabolic processes (Baker and Thummel, 2007; Chatterjee and Perrimon, 2021; Gáliková and Klepsatel, 2018; Ugur et al., 2016). *Drosophila* development usually takes around ten days, with the embryonic stage, three larval stages (L1, L2, and L3 instar larval stages), non-feeding wandering larval stage, and papal stage. Anabolism pathways, including DNL, are highly activated during larval growth. Therefore, *Drosophila* larvae can gain more than 200 folds of body mass in less than four days (Church and Robertson, 1966; Slaidina et al., 2009). In consist with that, triglycerides (TAG) storage increases significantly during larval development, especially at the L3 stage **(Figure 1A)**.

**Figure 1.**
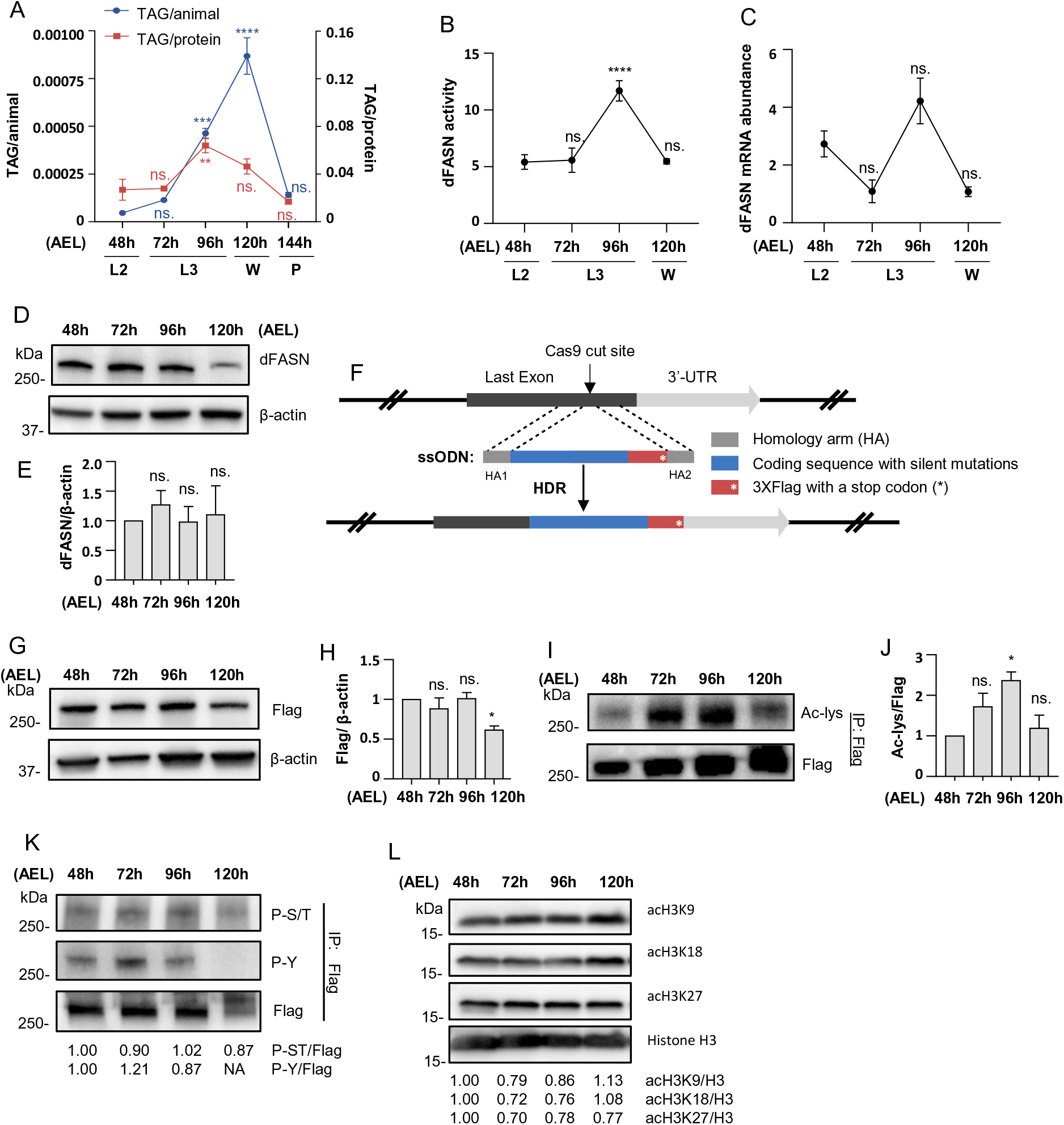
Acetylation modification of dFASN, but not protein expression, is positively correlated with DNL during *Drosophila* larval development. (A) Stage-specific triglyceride (TAG) level in *Drosophila* larvae. TAG was normalized against total protein amount or number of larvae (or pupa). One-way ANOVA (*vs.* 48h). Error bars represent SD; \*\*\*\**p*<0.0001; \*\*\**p*<0.001; \*\**p*<0.01; ns. *p*>0.05. AEL: after egg laying. L2: 2nd instar larvae. L3: 3rd instar larvae. W: wandering larvae. P: pupae. (B) Stage-specific dFASN enzymatic activity. dFASN activity was determined by oxidized NADPH per minute per mg of total protein. One-way ANOVA (*vs.* 48h). Error bars represent SD; \*\*\*\**p*<0.0001; ns. *p*>0.05. (C) Stage-specific dFASN mRNA abundance analyzed by qRT-PCR. One-way ANOVA (*vs.* 48h). Error bars represent SD; ns. *p*>0.05. (D) Stage-specific dFASN protein expression analyzed by western blotting with an antibody recognizing dFASN (anti-dFASN). (E) Quantification of dFASN protein expression (panel D) normalized against β-actin. Error bars represent SD. One-way ANOVA (*vs*. 48h), ns.: *p*>0.05. (F) Design of FASN-Flag knock-in fly line using CRISPR/Cas9-mediated homology-directed repair (HDR) and ssODNs (single-stranded oligodeoxynucleotides). The precise Cas9 cutting position was at the last exon adjacent to the stop codon. (G) Stage-specific endogenous FASN-Flag protein expression analyzed by western blotting with anti-Flag antibody. (H) Quantification of dFASN-Flag protein expression (panel G) normalized against β-actin. Error bars represent SD. One-way ANOVA (*vs*. 48h), \**p*<0.05, ns. *p*>0.05. (I) Stage-specific dFASN acetylation. dFASN-Flag was immunoprecipitated from *FASN^Flag^* larvae followed by western blotting analyses. dFASN acetylation was detected by anti-acetylated-lysine antibody (anti-Ac-lys). (J) Quantification of dFASN acetylation in panel I. Acetylated dFASN was normalized against total dFASN-Flag. Error bars represent SD. One-way ANOVA (*vs*. 48h). (K) Stage-specific dFASN phosphorylation. dFASN-Flag was immunoprecipitated from *FASN^Flag^* followed by western blotting analyses. dFASN phosphorylation was detected by anti-phospho-Ser/Thr (anti-P-S/T) and anti-phospho-Tyr (anti-P-Y) antibodies. Phospho-Ser/Thr and phospho-Tyr were normalized against dFASN-Flag expression, respectively. (L) Stage-specific Histone H3 acetylation analyzed by western blotting. Acetylation of H3K9, H3K18, and H3K27 were normalized against histone H3 expression, respectively.

Upregulation of DNL is a significant contributor to increased fat mass, which converts excess carbohydrates into fatty acids that are then esterified to storage TAG (Lin et al., 2005; Nagai et al., 2009). FASN is an essential lipogenic enzyme in DNL, synthesizing palmitate acids from acetyl-CoA, malonyl-CoA, and NADPH (Smith, 1994). To determine the role of *Drosophila* FASN (dFASN or FASN1) in TAG accumulation in growing larvae, we examined stage-specific dFASN enzymatic activity. We found that dFASN enzymatic activity was also upregulated with larval growth and was highly correlated with TAG levels **(Figure 1B)**. This result suggests that *de novo* fatty acid synthesis is vital for fat deposition in developing larvae.

It is known that the activation of FASN is mainly through upregulation of transcription and translation (Horton et al., 2003; Liang et al., 2002; Xu et al., 2013). To characterize how dFASN is regulated during larval development, we first determined the stage-specific mRNA and protein expressions of dFASN. Although the mRNA expression was slightly induced at L3 larval stages (96 hours after egg laying, 96 h AEL) **(Figure 1C)**, we surprisingly found that the protein expression of dFASN remained unchanged during larval development, from 48 to 96 hours AEL **(Figure 1D and 1E)**. These data suggest that transcriptional and translational regulation of dFASN may play a minor role in developmental lipogenesis, whereas PTMs might be involved in dFASN activation.

To investigate the PTMs of dFASN, we first generated a knock-in fly line through CRISPR/Cas9-mediated homology-directed repair (HDR) technique, in which a 3xFlag tag was precisely inserted at the 3’-end of the dFASN coding region **(Figure 1F)**. Three transformants containing the correct 3xFlag knock-in were recovered out of 230 G0 flies and verified by PCR, western blotting, and Sanger sequencing **(Figure S1A-D)**. The molecular weight of dFASN-3xFlag was around 270 kDa, close to the predicted molecular weight of dFASN **(Figure S1C)**. The knock-in lines are homozygous viable, fertile, and exhibit normal larval development (data not shown). Similar to the above western blot analysis using dFASN antibodies, the levels of endogenous dFASN proteins from 3xFlag knock-in flies also remained unchanged during larval development **(Figure 1G and 1H)**.

Phosphorylation and acetylation are two known modifications of FASN identified from previous studies (Hennigar et al., 1998; Jin et al., 2010; Lin et al., 2016). To characterize these two modifications during larval development, we immunoprecipitated endogenous dFASN from the knock-in fly line *FASN^Flag^* with an anti-Flag antibody followed by western blot analysis. Intriguingly, the acetylation levels of dFASN were highly correlated with its enzymatic activity, with a peak at 96 hours AEL **(Figure 1I and 1J)**. In comparison, phosphorylation of dFASN did not significantly change from 48 to 96 hours AEL **(Figure 1K)**. To exclude the possibility that increased dFASN acetylation at L3 larvae is due to global changes in acetylome, we measured three histone acetylation marks, acH3K9, acH3K18, and acH3K27, in developing larvae. Interestingly, none of the histone acetylation marks changed significantly throughout larval development **(Figure 1L)**, suggesting that upregulated dFASN acetylation is not due to changes in global acetylation. Taken together, our results indicate that acetylation modification of dFASN, instead of transcription and translation, is the major regulatory mechanism for DNL during *Drosophila* development.

### K813 is a crucial acetylated lysine for DNL, body fat accumulation, and normal *Drosophila* development

To identify the acetylated lysine sites of dFASN, we carried out a proteomic analysis with immunoprecipitated dFASN from *FASN^Flag^* Flies. Eight lysine residues that distribute on four different domains of dFASN were identified **(Figure 2A)**. When comparing these lysine sites with previous acetylome studies, we found four lysine residues, K813, K926, K1800, and K2466, that are conserved from flies to mammals and have been frequently acetylated in many animal species **(Figure 2B-2C)** (Beli et al., 2012; Bouchut et al., 2015; Chen et al., 2012; Choudhary et al., 2009; Elia et al., 2015; Hebert et al., 2013; Lundby et al., 2012; Mertins et al., 2013; Park et al., 2013; Shaw et al., 2011; Simon et al., 2012; Sol et al., 2012; Still et al., 2013; Svinkina et al., 2015; Weinert et al., 2013; Xu et al., 2016; Yang et al., 2011). Among these four lysine residues, K813 was the most frequently acetylated one, followed by K926 **(Figure 2B)**. Therefore, K813 and K926 were selected for further functional studies.

**Figure 2.**
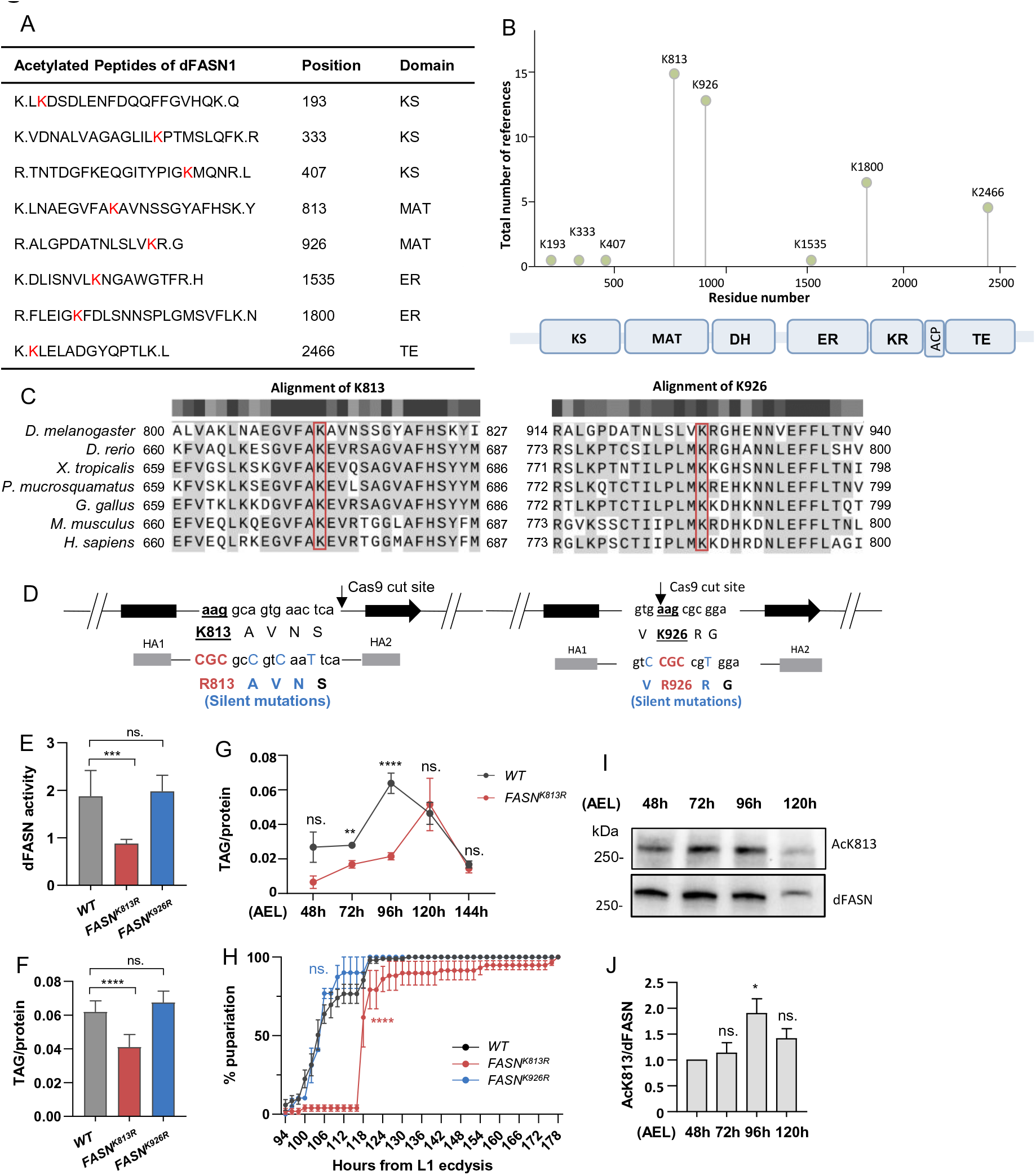
K813 is a crucial acetylated lysine for DNL, body fat accumulation, and normal *Drosophila* development. (A) Table summarizing mass-spectrometry based analysis of dFASN acetylation. Acetylated lysine residues are in red color. Residue number of the acetylated site and the associated domain are indicated. (B) The distribution and detection frequency of acetylated lysine residues of mammal and fly FASN reported in the literature. The number of records in which this acetylation site was assigned using proteomic discovery mass-spectrometry is shown on y-axis. The residue number of the acetylated site and its localization on FASN domain is shown on x-axis. (C) Sequence alignment of lysine residues K813 and K926 among species. *Drosophila melanogaster, D. melanogaster; Danio rerio, D. rerio; Xenopus tropicalis, X. tropicalis; Protobothrops mucrosquamatus, P. mucrosquamatus; Gallus gallus, G. gallus; Mus musculus, M. musculus; Homo sapiens, H. sapiens*. (D) Design of acetylation deficient fly mutants (*FASN^K813R^* and *FASN^K926R^*) using CRISPR/Cas9-mediated HDR. Sequence of ssODNs are shown. (E) dFASN enzymatic activity of *WT* (*yw^R^*), *FASN^K813R^*, and *FASN^K926R^* at L3 stage. dFASN activity was determined by oxidized NADPH per minute per mg of protein. One-way ANOVA (*vs. yw^R^*). Error bars represent SD; \*\*\**p*<0.001; ns. *p*>0.05. (F) TAG level of *WT* (*yw^R^*), *FASN^K813R^*, and *FASN^K926R^* at L3 stage. TAG was normalized against total protein amount. One-way ANOVA (*vs. yw^R^*). Error bars represent SD; \*\*\*\**p*<0.0001; ns. *p*>0.05. (G) Stage-specific TAG level of *WT* (*yw^R^*) and *FASN^K813R^*. Multiple *t*-test. Error bars represent SD; \*\**p*<0.01; \*\*\*\**p*<0.0001; ns. *p*>0.05. (H) Developmental timing of *WT* (*yw^R^*), *FASN^K813R^*, and *FASN^K926R^*. Number of pupariation was counted every 2-4 hours. Log-rank test (*vs. yw^R^*). Error bars represent SD; \*\*\*\**p*<0.0001; ns. *p*>0.05. (I) Site-specific acetylation at K813 in larval development stages determined by western blotting analyses. Acetylation of K813 was measured by an antibody specifically recognizing acetylated K813 (anti-AcK813). (J) Quantification of K813 acetylation in panel I. K813 acetylation was normalized against total dFASN. One-way ANOVA (*vs*. 48h). Error bars represent SD. \**p*<0.05; ns. *p*>0.05.

To study the functional role of the identified lysine residues in dFASN enzymatic activity, lipogenesis, and larval development, we generated two acetylation-deficient mutants (Lys to Arg substitution), *FASNK^813R^* and *FASNK^926R^*, through CRISPR/Cas9-mediated HDR **(Figure 2D)**. Successful transformants were verified by PCR using primers targeting mutation regions and Sanger sequencing **(Figure S2A-S2C)**. Two transformants that contained K813>R813 substitution and one transformant that contained K926>R926 substitution were recovered **(Figure S2A)**. *FASN^K813R^* are homozygous viable and fertile, while *FASN^K926R^* are homozygous viable and homozygous infertile **(Figure S2D)**.

To determine whether lysine acetylation potentially mediates the function of dFASN, we measured dFASN activity in *FASN^K813R^* and *FASN^K926R^* at 96 hours AEL. Remarkably, *FASN^K813R^* mutants, but not *FASN^K926R^*, showed decreased dFASN enzymatic activity **(Figure 2E)**. Body fat accumulation at 96 hours AEL of the two mutants was also determined. As was shown in **Figure 2F,** the amount of TAG was significantly reduced in *FASN^K813R^*, but not in *FASN^K926R^*. When examining the TAG levels throughout developmental stages, we found that *FASN^K813R^* mutants exhibited lower TAG levels at almost all stages, with a delayed peak of TAG at around 120 hours AEL **(Figure 2G)**.

Dysregulated fatty acid biosynthesis impairs larval growth and development (Xie et al., 2015). Consistent with this idea, we found that pupariation of *FASN^K813R^* mutants was significantly delayed, while *FASN^K926R^* exhibited normal development **(Figure 2H)**. Sufficient larval-derived body fat is required for normal development of adult tissue at pupal stage, and insufficient lipid accumulation in larval development may cause growth deficiency (Aguila et al., 2007). Indeed, the body weight of *FASN^K813R^* adults (both male and female) was significantly decreased (**Figure S2E**).

Given that the acetylation of K813, not K926, is required for the activation of dFASN activity, increased fat accumulation, and normal larval development, we speculate that K813 acetylation may be positively correlated with dFASN activity and peak at 96 hours AEL (L3 larval stage). To test this idea, we measured acetylation of K813 at different larval stages with a newly generated antibody specific to acetylated K813 (acK813). The specificity of our acK813 antibody was verified through dot plot analysis, which confirmed that the acK813 antibody showed cross-reactivity with only acetylated peptide, not non-acetylated peptide (**Figure S3A**). As predicted, the acetylation of K813 significantly increased at 96 hours AEL **(Figure 2I-2J)**, which is positively correlated with dFASN activity **(Figure 1B)**.

Taken together, our studies demonstrate that disturbance of site-specific acetylation at K813 decreases dFASN activity, slows down lipid accumulation and animal development, indicating that K813 is a critical acetylated lysine site in regulating lipid synthesis and *Drosophila* development. Interestingly, although dFASN activity is impaired in *FASN^K813R^*, enzyme function was not destroyed, and the mutants can still develop into viable pupae and adults. While on contrast, dFASN knockout mutants are homozygous lethal (**Figure S2D**). Therefore, we hypothesize that acetylation at K813 fine-tunes dFASN activity and lipogenesis during larval development.

### Acetylation at K813 increases dFASN activity at high substrate concentrations

Eukaryotic FASN is a multifunctional protein containing seven activity domains (Leibundgut et al., 2008). K813 is localized at the malonyl/acetyltransferase (MAT) domain **(Figure 2A-2B)**, which is responsible for binding substrates acetyl-CoA (initiation primer) and malonyl-CoA (chain extender) (Smith, 1994). To further investigate how acetylation at K813 modulates dFASN activity, we first examined the crystal structure of MAT domains of both dFASN and human FASN (hFASN) **(Figure 3A)** (Pappenberger et al., 2010). Interestingly, K673 of hFASN (homolog of dFASN K813) was at the substrate docking pocket of the MAT domain. Since the crystal structure of dFASN is not currently available, we predicted the protein structure by the I-TASSER server (Roy et al., 2010; Yang et al., 2015; Zhang, 2008). The predicted structure *of Drosophila* protein revealed a similar localization of K813 as that of hFASN K673 **(Figure 3A)**.

**Figure 3.**
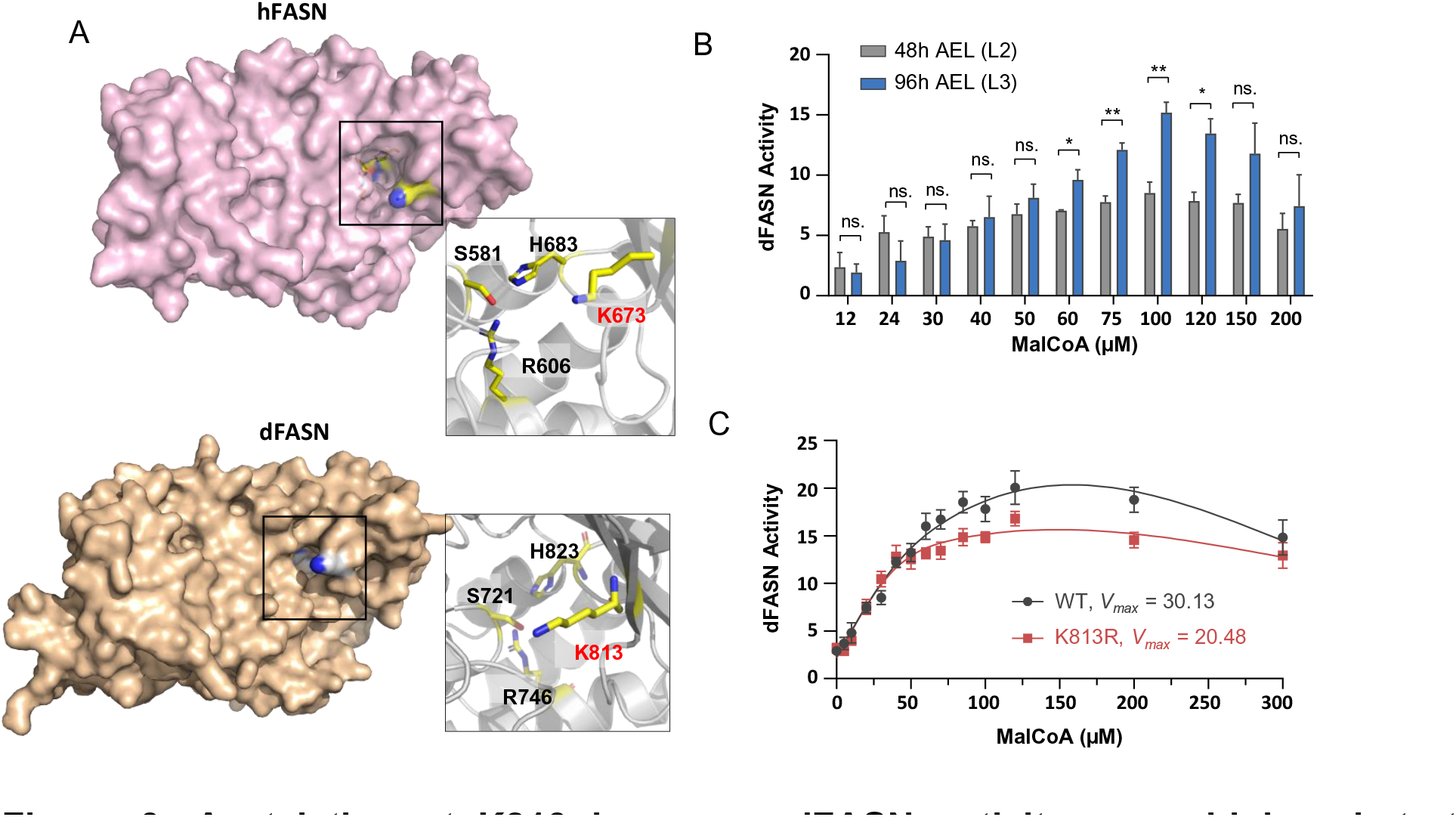
Acetylation at K813 increases dFASN activity upon high substrate concentrations. (A) The structure of MAT domain human FASN (hFASN) (PDB: 3hhd) (Pappenberger *et al*., 2010) (up) and predicted MAT domain structure of dFASN (down). The boxes indicated the substrate docking pocket of the MAT domain. The insets showing the acetylated lysine residue (K673 in human and K813 in fly) and the key residues of the active sites of the MAT domain. The protein structures were visualized with PyMOL Molecular Graphics System. (B) Malonyl-CoA (MalCoA)-dependent dFASN activity at L2 and L3. *t*-test. Error bars represent SD. \**p*<0.05; \*\**p*<0.01; ns. *p*>0.05. (C) Steady-state kinetics evaluation of WT and K813R recombinant dFASN upon malonyl-CoA. Error bars represent SD. Kinetics parameter *V_max_* was calculated according to Michaelis-Menten analysis.

The proximity of K813 to the substrate docking pocket of the MAT domain hints us to test whether acetylation of K813 modulates dFASN catalytic activity upon malonyl-CoA, the main substrates of MAT domain. To test this idea, we first measured the activity of dFASN from 48 and 96 hours AEL larvae by incubating protein lysates with different concentrations of substrate malonyl-CoA **(Figure 3B)**. Although dFASN activity of both groups increased upon high malonyl-CoA concentration, dFASN from 96 hours AEL is more sensitive to the change of malonyl-CoA levels. The activity of dFASN from the two larval stages was similar at low malonyl-CoA concentrations (<60 µM), whereas dFASN activity from 96 hours AEL larvae was significantly higher than that of 48 hours AEL upon high malonyl-CoA concentrations (60-120 µM) **(Figure 3B)**. Additionally, substrate-excess inhibition was observed under high malonyl-CoA dosages (120-200 µM) in both groups, consistent with previous kinetics studies on yeast FAS (Singh et al., 2020). Since dFASN protein expressions were similar between the two stages, we speculated that the alteration in malonyl-CoA-dependent dFASN activity may be caused by acetylation.

To directly investigate the role of K813 acetylation in dFASN enzymatic kinetics upon malonyl-CoA, we expressed recombinant dFASN proteins, both wild-type (WT) and K813R mutant, using Bac-to-Bac expression system followed by His-resin purification. Purified recombinant proteins were verified by Coomassie Blue staining **(Figure S3B)**, which were then used for steady-state kinetics evaluation. The recombinant dFASN proteins with K813R substitution showed ∼30% reduction of the maximum rate of reaction (*V_max, Malonyl-CoA_*) compared to WT proteins **(Figure 3C)**. Consistent with dFASN activity in L3 larvae, WT recombinant proteins exhibited higher activity than K813R mutants upon higher malonyl-CoA concentrations (50-200 µM), but not upon low malonyl-CoA concentrations (<50 µM) **(Figure 3C)**. These results showed that interfering with the acetylation of K813 alters the kinetics properties of dFASN.

Taken together, our findings show that K813 localizes at the substrate docking pocket of the MAT domain, and acetylation of K813 may alter the conformation of the docking pocket to enhance enzyme activity at high substrate concentrations. Here, we propose a fine-tune mechanism of dFASN regulation in which acetylation of K813 modulates dFASN catalytic activity in response to fluctuated substrate availability during larval development.

### K813 is not acetylated through known *Drosophila* KATs

The unique pattern of dFASN acetylation indicates that it is tightly regulated during *Drosophila* development. We next asked how acetylation of dFASN, especially K813, is regulated. Like histone acetylation, non-histone proteins are primarily acetylated and deacetylated by acetyltransferases (KATs) and deacetylases (KDACs), respectively (Narita et al., 2019). To date, eight KATs with different subcellular localizations have been identified in the *Drosophila* genome **(Figure 4A)**. To identify the KAT that regulates dFASN acetylation in developing larvae, we first checked the mRNA expression of *Drosophila* KATs in previous transcriptomic analysis (Graveley et al., 2011). Interestingly, none of the KATs showed upregulation in L3 stage when dFASN acetylation is elevated **(Figure 4B)**.

**Figure 4.**
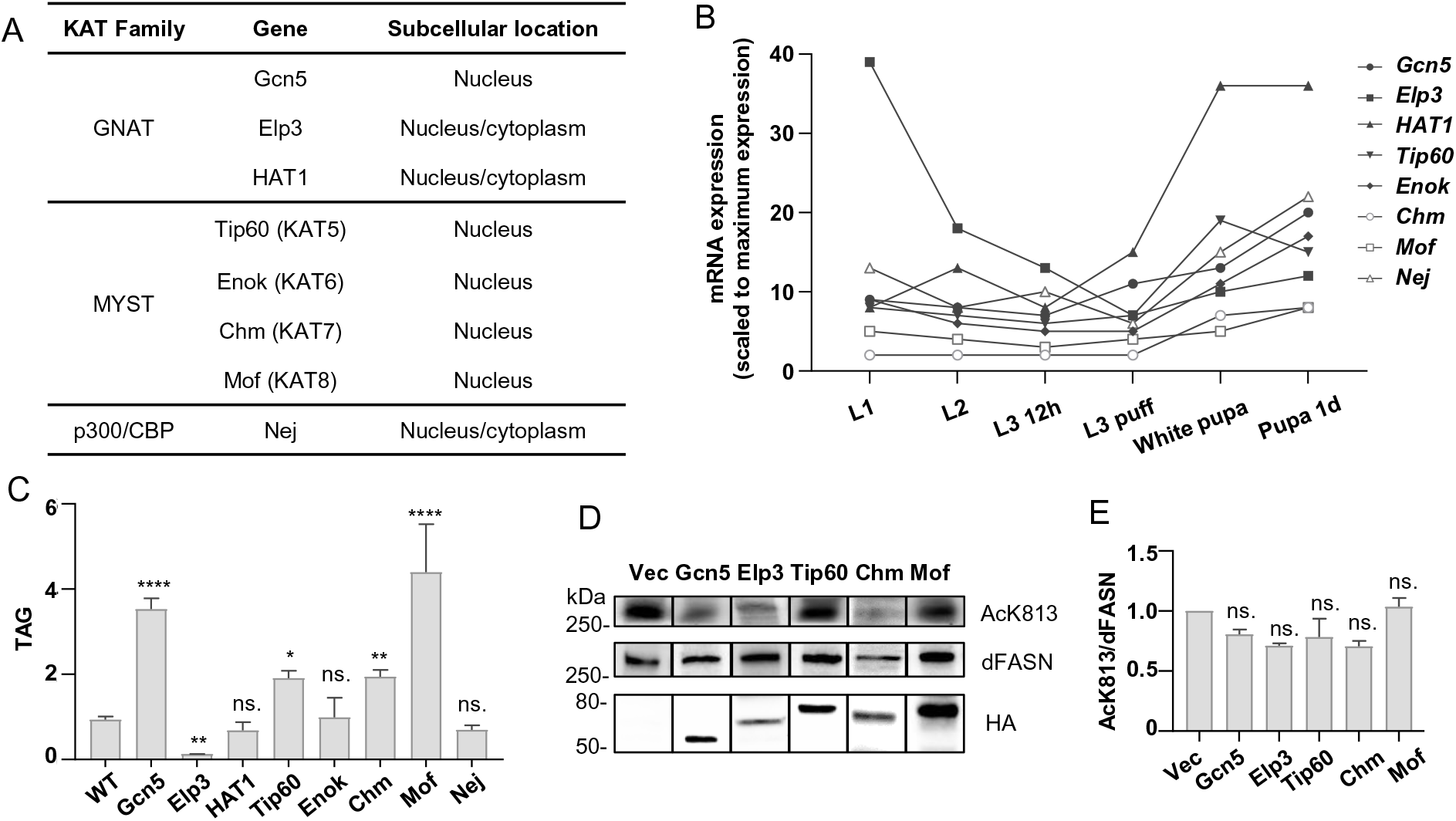
K813 is not acetylated through known *Drosophila* KATs. (A) Table summarizing known *Drosophila* KATs and their subcellular localizations. (B) Developmental mRNA expression of KATs. The transcriptional profiles of KATs were retrieved from the *Drosophila* developmental transcriptome study (Graveley, Brooks et al. 2011). (C) TAG levels of KAT mutants at L3 larval stage. TAG was normalized against total protein. Y-axis represents folds change *vs.* WT (*yw^R^ or w^1118^*). One-way ANOVA (*vs. WT*). Error bars represent SD; \*\*\*\**p*<0.0001; \*\**p*<0.01; \**p*<0.05; ns. *p*>0.05. (D) Acetylation of K813 under overexpression of KATs in *Drosophila* Kc167 cells determined by western blotting analyses with anti-AcK813. (E) Quantification of K813 acetylation under overexpression of KATs in panel D. K813 acetylation was normalized against total dFASN. Error bars represent SD. One-way ANOVA (*vs*. control). ns. *p*>0.05.

To further investigate the role of *Drosophila* KATs in dFASN acetylation and lipogenesis, we performed a genetic screen to identify the KATs involved in TAG accumulation. Interestingly, loss-of-function mutations of four *Drosophila* KATs (Gcn5, Tip60, Chm, and Mof) exhibited elevated TAG levels, while Elp3 mutants showed reduced TAG at L3 larval stage **(Figure 4C)**. These findings suggest that Epl3 may be a positive regulator for lipogenesis and potentially dFASN acetylation.

Lastly, we directly examined the role of *Drosophila* KATs in dFASN acetylation by overexpressing the KATs in *Drosophila* Kc167 cells and measured the acetylation level of K813 using acK813 antibodies. Five KATs that were shown to regulate TAG accumulation were cloned to expression vectors. Unexpectedly, our western blot results showed that ectopic expression of all five KATs (including Elp3) failed to induce acetylation of K813 **(Figure 4D-4E)**. Thus, these data suggest that K813 might be acetylated by an unknown KAT, or the acetylation of K813 is regulated through a unique KAT-independent mechanism.

### K813 is rapidly autoacetylated by acetyl-CoA in a dosage-dependent manner

Except being enzymatically catalyzed by KATs, lysine acetylation also arises from a nonenzymatic reaction with acetyl-CoA, also known as chemical acetylation or autoacetylation. Nonenzymatic acetylation favors alkaline environments (pH=8.0) with high acetyl-CoA concentrations (0.1-1.5 mM), like mitochondrial matrix (Hansen et al., 2019; Olia et al., 2015; Wagner and Payne, 2013). In contrast, nonenzymatic acetylation of cytosolic proteins is rarely observed due to low acetyl-CoA concentrations (3-30 µM) and neutral pH environment (Olia et al., 2015). Interestingly, nonenzymatic acylation of FASN was observed in a recent global nonenzymatic acylation screen (Kulkarni et al., 2017). However, no follow-up validation was performed to confirm if cytosolic protein FASN is nonenzymatically modified by either acetyl-CoA or other acyl-CoAs.

To explore the possibility that dFASN acetylation is regulated through a nonenzymatic mechanism, we measured the acetylation of dFASN *in vitro* by incubating recombinant dFASN with 5-20 µM of acetyl-CoA, the dosages close to actual cytosolic acetyl-CoA levels (Wagner and Payne, 2013). Intriguingly, recombinant dFASN was rapidly acetylated by as few as 5 µM of acetyl-CoA at both pH7.0 and pH8.0 in one hour **(Figure 5A-5B)**. Consistent with the idea that nonenzymatic acetylation favors alkaline conditions (Baldensperger and Glomb, 2021; Wagner and Payne, 2013), the acetylation levels of recombinant dFASN were higher at pH8.0 than at pH7.0. More importantly, our observations on dFASN acetylation at pH7.0 upon low concentrations of acetyl-CoA suggest that endogenous dFASN proteins could be autoacetylated *in vivo*.

**Figure 5.**
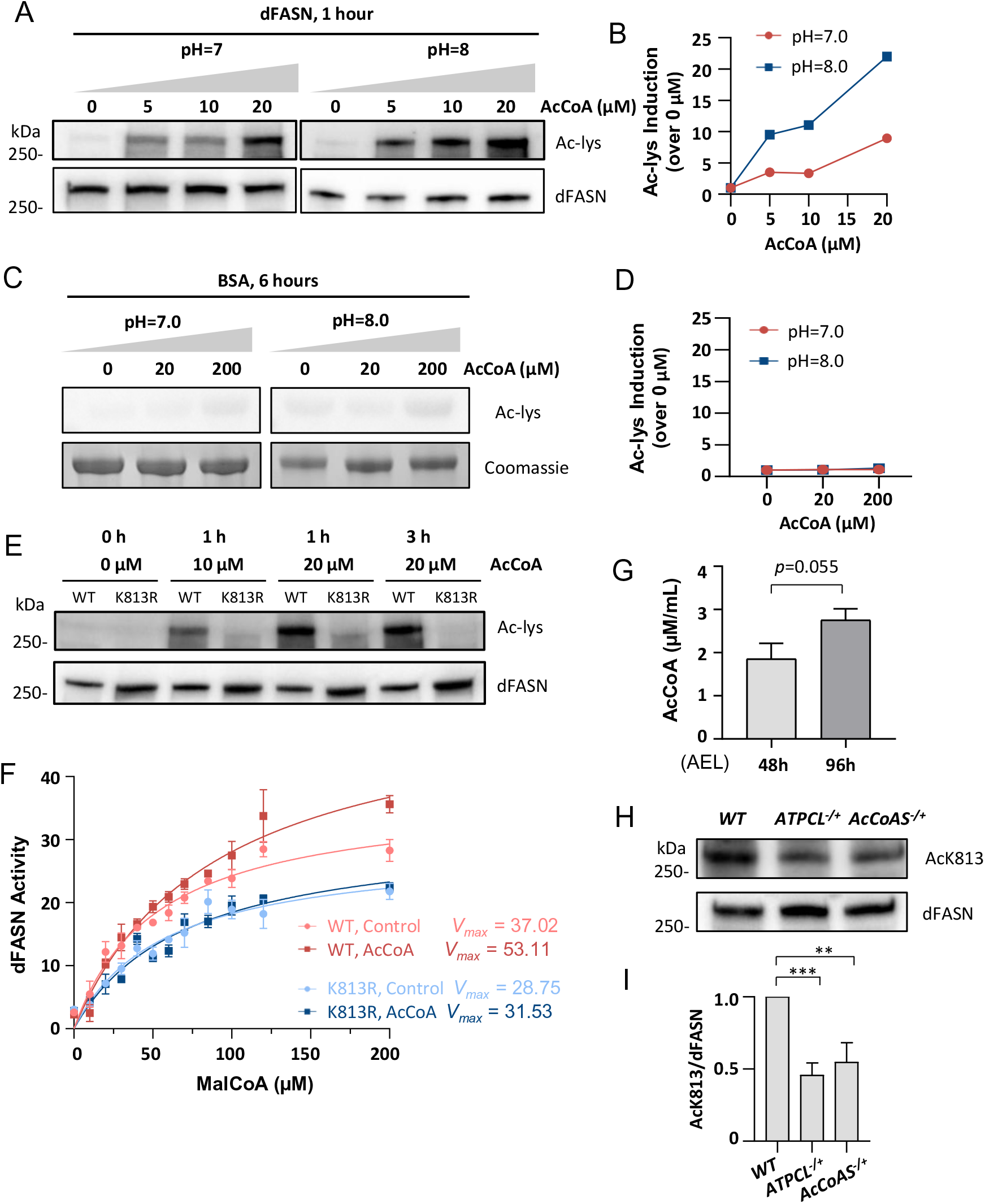
K813 is rapidly autoacetylated by acetyl-CoA in a dosage-dependent manner. (A-B) Acetylation of dFASN by acetyl-CoA (AcCoA). Recombinant dFASN was incubated with 0, 5, 10, or 20 µM of AcCoA at pH7.0 or pH8.0 for one hour prior to western blotting analyses with anti-Ac-lys. Folds of induction over 0 µM of AcCoA treatment were shown in (B). (C-D) Acetylation of BSA by AcCoA treatment. BSA was incubated with 0, 20, or 200 µM of AcCoA at pH7.0 or pH8.0 for six hours prior to western blotting analyses with anti-Ac-lys. Folds of induction over 0 µM of AcCoA treatment were shown in (D). (E) Acetylation of WT or K813R dFASN by AcCoA treatment. WT or K813R recombinant dFASN was incubated with 0, 10, or 20 µM of AcCoA at pH8.0 for one hour or three hours prior to western blotting analyses. (F) Steady-state kinetics evaluation of WT and K813R of recombinant dFASN post AcCoA treatment. WT or K813R recombinant dFASN were incubated with 20 µM of AcCoA at pH7.0 for 0 or 1 hour before activity assay. Error bars represent SD. Kinetics parameter *V_max_* was calculated according to Michaelis-Menten analysis. (G) Quantification of intracellular AcCoA levels of developing *Drosophila* larvae. L2: 48h AEL; L3: 96h AEL. One tail *t*-test. Error bars represent SD. (H) Acetylation of K813 in *WT* (*yw^R^*), *ATPCL^-/+^*, and *AcCoAS^-/+^* at L3 larval stage. (I) Quantification of K813 acetylation in in panel H. K813 acetylation was normalized against total dFASN. Error bars represent SD. One-way ANOVA (*vs. WT).* \*\*\**p*<0.001; \*\**p*<0.01.

In contrast, the negative control protein Bovine Serum Albumin (BSA) showed no autoacetylation, even when the proteins were incubated with a relatively high concentration of acetyl-CoA (200 µM) for six hours **(Figure 5C-5D)**. More interestingly, autoacetylation of dFASN was largely blocked by K813R substitution **(Figure 5E)**, suggesting that K813 is the primary residue of dFASN that could be autoacetylated.

Given that dFASN can be acetylated by acetyl-CoA in a dosage-dependent manner **(Figure 5A-5B)**, an autoregulatory mechanism might be used by developing larvae to fine-tune dFASN acetylation and enzymatic activity in response to fluctuating cytosolic acetyl-CoA levels. We then asked whether acetyl-CoA-mediated autoacetylation of K813 modulates dFASN activity. To test this idea, we evaluated steady-state kinetics of recombinant dFASN protein incubated with or without 20 µM of acetyl-CoA. Intriguingly, recombinant dFASN proteins pre-incubated with acetyl-CoA showed 43% increases in the maximum rate of reaction *V_max, Malonyl-CoA_* **(Figure 5F)**. Importantly, *V_max, Malonyl-CoA_* of K813R mutant protein was not altered by acetyl-CoA treatment **(Figure 5F)**.

Taken together, our findings demonstrate that cytosolic protein dFASN could be rapidly autoacetylated by acetyl-CoA in a dosage-dependent manner at neutral conditions, and K813 is the major autoacetylation site. The autoacetylation of K813 by acetyl-CoA plays an important role in fine-tuning dFASN enzymatic activity.

### Acetylation of K813 is regulated by ATPCL and AcCoAS

To explore whether K813 acetylation is regulated by the acetyl-CoA flux *in vivo*, we first measured cellular acetyl-CoA of 48 and 96 hours AEL larvae by LC-MS/MS. Interestingly, the levels of acetyl-CoA increased at 96 hours AEL compared to 48 hours AEL **(Figure 5G)**, which is positively correlated with dFASN acetylation and developmental lipogenesis. ATP citrate lyase (ATPCL) and acetyl coenzyme A synthase (AcCoAS) are two main contributors to the cytosolic acetyl-CoA pool, producing acetyl-CoA from citrate and acetate, respectively (Pietrocola et al., 2015). Interestingly, mRNA expressions of the two genes are both upregulated at L3 sage **(Figure S3C)** (Graveley et al., 2011). This finding is consistent with our LC-MS/MS results showing elevated acetyl-CoA levels in fast-growing L3 larvae.

Furthermore, to directly determine whether ATPCL or AcCoAS regulates dFASN acetylation *in vivo*, we measured acetylation of K813 in ATPCL and AcCoAS loss-of-function mutants. Our western blot analysis showed that K813 acetylation was reduced 50% in both *ATPCL^[01466]/+^ and AcCoAS^[MI12066]/+^* loss-of-function mutants at 96 hours AEL **(Figure 5H-5I)**, suggesting that acetylation of K813 is sensitive to cytosolic acetyl-CoA flux. Taken together, our results indicate that acetylation of dFASN is regulated by both ATPCL and AcCoAS, two major enzymes controlling cytosolic acetyl-CoA biosynthesis. Autoacetylation of K813 is likely mediated by intracellular acetyl-CoA flux *in vivo*.

### N-xx-G-x-A motif is required for autoacetylation of K813

The mechanism of nonenzymatic acetylation, especially of cytosolic proteins, is poorly understood. It has been shown that mitochondrial proteins can be acetylated when incubated with high dosages of acetyl-CoA (200 µM to 1.5 mM) in alkaline buffer (pH=8.0) for over three hours (James et al., 2017; Wagner and Payne, 2013). However, our results revealed that dFASN could be acetylated under a much tougher condition (5-20 µM of acetyl-CoA and neutral pH), which suggests that an efficient and unique mechanism is involved in acetyl-CoA-mediated dFASN autoacetylation.

To uncover the mechanism underlying dFASN autoacetylation, we expressed recombinant KS-MAT didomain of dFASN using in *E. coli* BL21 (DE3) expression system. KS-MAT didomain, but not full-length dFASN, was chosen is because full-length dFASN (∼270 kDa) expressed from *E. coli* system may not fold properly. Functional KS-MAT has been successfully expressed from *E. coli* in previous studies (Rittner et al., 2018). The purified recombinant protein was verified by Coomassie Blue staining **(Figure S3D)**. To validate the autoacetylation of KS-MAT recombinant protein, we incubated recombinant KS-MAT with different amounts of acetyl-CoA for 1 hour and checked the acetylation of K813 by western blot analysis. Consistent with dFASN recombinant proteins produced by Bac-to-Bac expression system, recombinant KS-MAT was rapidly autoacetylated by acetyl-CoA in a dosage-dependent manner **(Figure 6A)**, although the induction of autoacetylation was not as strong as the full-length dFASN **(Figure 5A)**. Acetylation of recombinant KS-MAT under 20 µM acetyl-CoA treatment was not prominent, which could be due to the differences in protein folding between prokaryotic and eukaryotic expression systems. Therefore, 200 µM of acetyl-CoA was applied for the following analyses. We further confirmed that K813R substitution blocked acetyl-CoA-mediated autoacetylation of recombinant KS-MAT proteins **(Figure 6B)**. In contrast, K926R substitution did not affect autoacetylation of KS-MAT by acetyl-CoA treatment **(Figure 6B)**.

**Figure 6.**
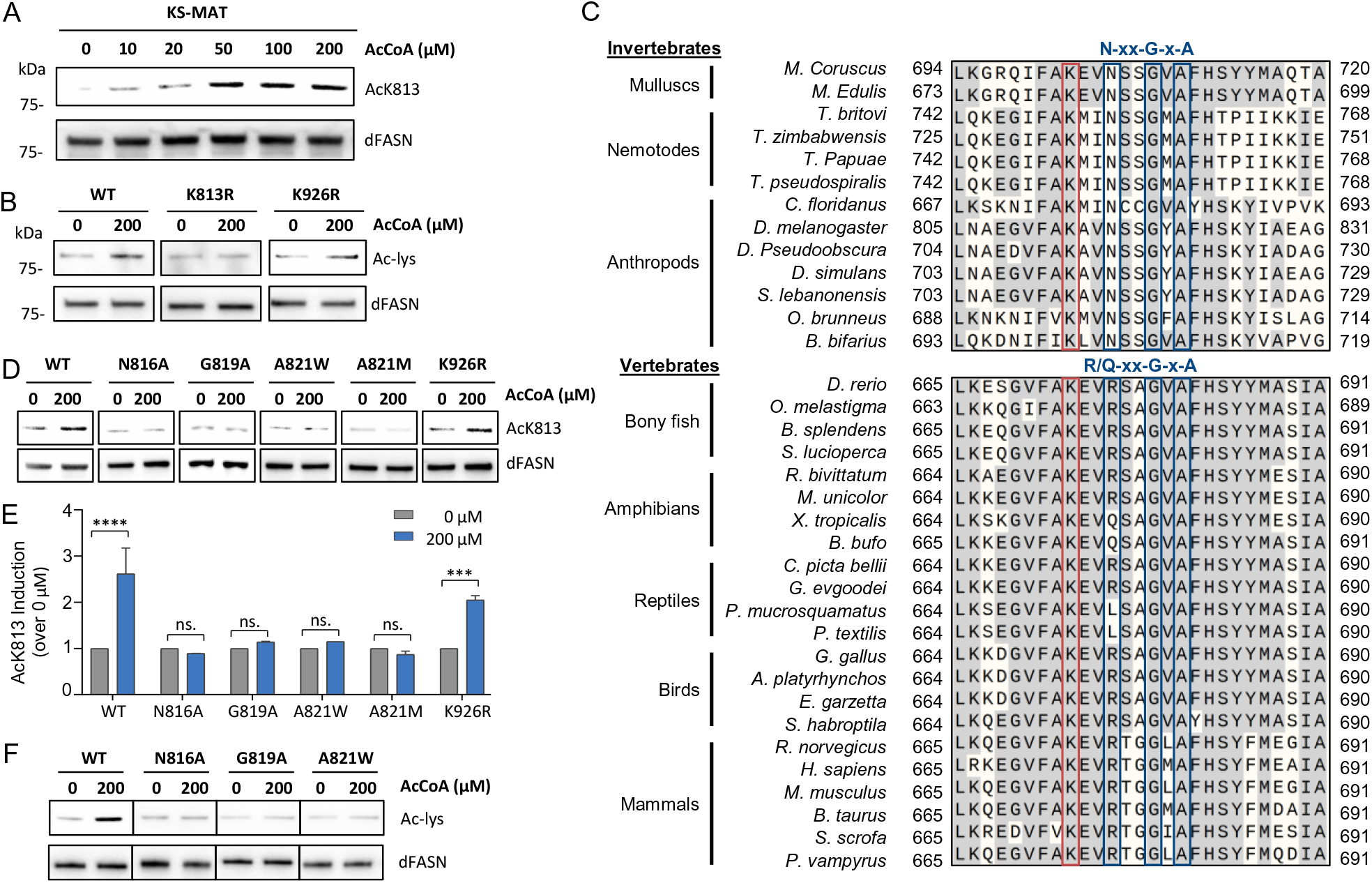
N-xx-G-x-A motif is required for autoacetylation of K813. (A) K813 acetylation of KS-MAT by AcCoA treatment. Recombinant KS-MAT didomain was incubated with 0, 10, 20, 50, 100, and 200 µM of AcCoA at pH8.0 for one hour or three hours prior to western blotting analyses with anti-AcK813. (B) Total acetylation of WT and mutant KS-MAT by AcCoA treatment. WT, K813R, or K926R recombinant KS-MAT was incubated with 200 µM of AcCoA at pH8.0 for one hour prior to western blotting analyses with anti-Ac-lys. (C) Sequence alignment of MAT domain (the region surrounding acetylated K813) among invertebrate and vertebrate species. Acetylated lysine is highlighted in red. The P-loop-like motif (N-xx-G-x-A in invertebrates or R/Q-xx-G-x-A in vertebrates) is highlighted in blue. (D) AcCoA-mediated K813 acetylation of single-site mutations (N816A. G819A, A821W, A821W, and K926R). WT or mutant KS-MAT recombinant proteins was incubated with 200 µM of AcCoA at pH8.0 for one hour prior to western blotting analyses with anti-AcK813. (E) Quantification of K813 acetylation of single-site mutations in panel D. K813 acetylation was normalized against total dFASN. (F) Total acetylation of single-site mutations by AcCoA treatment. WT, N816A, G819A, or A821W recombinant KS-MAT was incubated with 200 µM of AcCoA at pH8.0 for one hour prior to western blotting analyses with anti-Ac-lys.

Previous studies found that high stoichiometry acetylation of lysine is associated with neighboring cysteine (James et al., 2018), and peptides contained a cysteine near a lysine residue exhibit increased nonenzymatic acetylation *in vitro* (James et al., 2017). However, we could not locate any cysteine near K813 residue from the primary and tertiary structure of dFASN. Since dFASN can be rapidly acetylated by low concentration of acetyl-CoA at the neutral condition, we wonder whether dFASN has adapted some features of KATs to facility its autoacetylation. Intriguingly, we found a highly conserved motif (N-xx-G-x-A, where x denotes any amino acid), two amino acids away from K813, which resembles the signature P-loop sequence (R/Q-xx-G-x-A/G) of KATs (Roth et al., 2001). The alignment of the FASN amino acid sequences from a variety of animal species revealed that the first position of the motif is conditionally conserved (N in invertebrates and R/Q in vertebrates), while the fourth (G) and sixth (A) positions of the motif are highly conserved across all species **(Figure 6C)**.

P-loop is an invariant sequence in KATs motif A for acetyl-CoA recognition and binding (Dyda et al., 2000; Wolf et al., 1998). The three conserved residues form hydrogen bonds with pyrophosphate of acetyl-CoA (Wolf et al., 1998). KATs activity is largely disrupted when mutating any of the three signature residues (Kuo et al., 1998; Wang et al., 1998). To investigate the possible role of the N-xx-G-x-A motif in K813 autoacetylation, we expressed three recombinant proteins carrying N816A, G819A, or A821W substitutions using *E. coli* expression system (**Figure S3D**). Wild-type and mutant proteins were incubated with 200 µM of acetyl-CoA for one hour, followed by western blot analysis **(Figure 6D-6E)**. Strikingly, the induction of K813 acetylation was blocked by all single-site mutations, N816A, G819A, and A821W. Since A821 is not a surface residue (data not shown), to exclude the possibility that the blockage of autoacetylation by A821W mutant is due to the disruption of protein structure, we also tested another substitution, A821M. Similarly, recombinant proteins with A821M substitution showed no induction of K813 autoacetylation by acetyl-CoA treatment **(Figure 6D-6E)**. In addition, these single-site substitution mutations, N816A, G819A, and A821W, blocked total acetylation of KS-MAT recombinant proteins (using Ac-lys antibody), which further suggests that K813 is the major lysine residue that is autoacetylated **(Figure 6F)**. Together, these results demonstrate that N-xx-G-x-A motif plays a vital role in the autoacetylation of K813. It is likely that N-xx-G-x-A motif of dFASN is responsible for acetyl-CoA recognition and binding, similar to the P-loop function in KATs. Future experiments are needed to examine this possibility.

### Sirt1-mediated deacetylation of dFASN regulates lipogenesis and developmental timing in *Drosophila* larvae

Finally, we examined how K813 is deacetylated. There are ten KDACs in the *Drosophila* genome with distinct subcellular localizations, five from the HDACs family and five from the SIRTs family **(Figure 7A)**. When examining the developmental expression profiles of these KDACs and SIRTs (Graveley et al., 2011), we found that mRNA expression of *Sirt1* is high at early larval development (L1) and pupa stage, while it is reduced at L3 larval stage **(Figure 7B)**. The expression pattern of *Sirt1* negatively correlates with dFASN acetylation, suggesting Sirt1 might be the KDAC deacetylating dFASN. Furthermore, we conducted a genetic screen to examine the role of KDACs in TAG accumulation. Loss-of-function mutations of three sirtuins (Sirt1, Sirt4, and Sirt7) showed elevated TAG levels, suggesting a negative role of these sirtuins in lipogenesis and potentially dFASN acetylation **(Figure 7C)**. However, only Sirt1 is localized in the cytoplasm **(Figure 7A)**.

**Figure 7.**
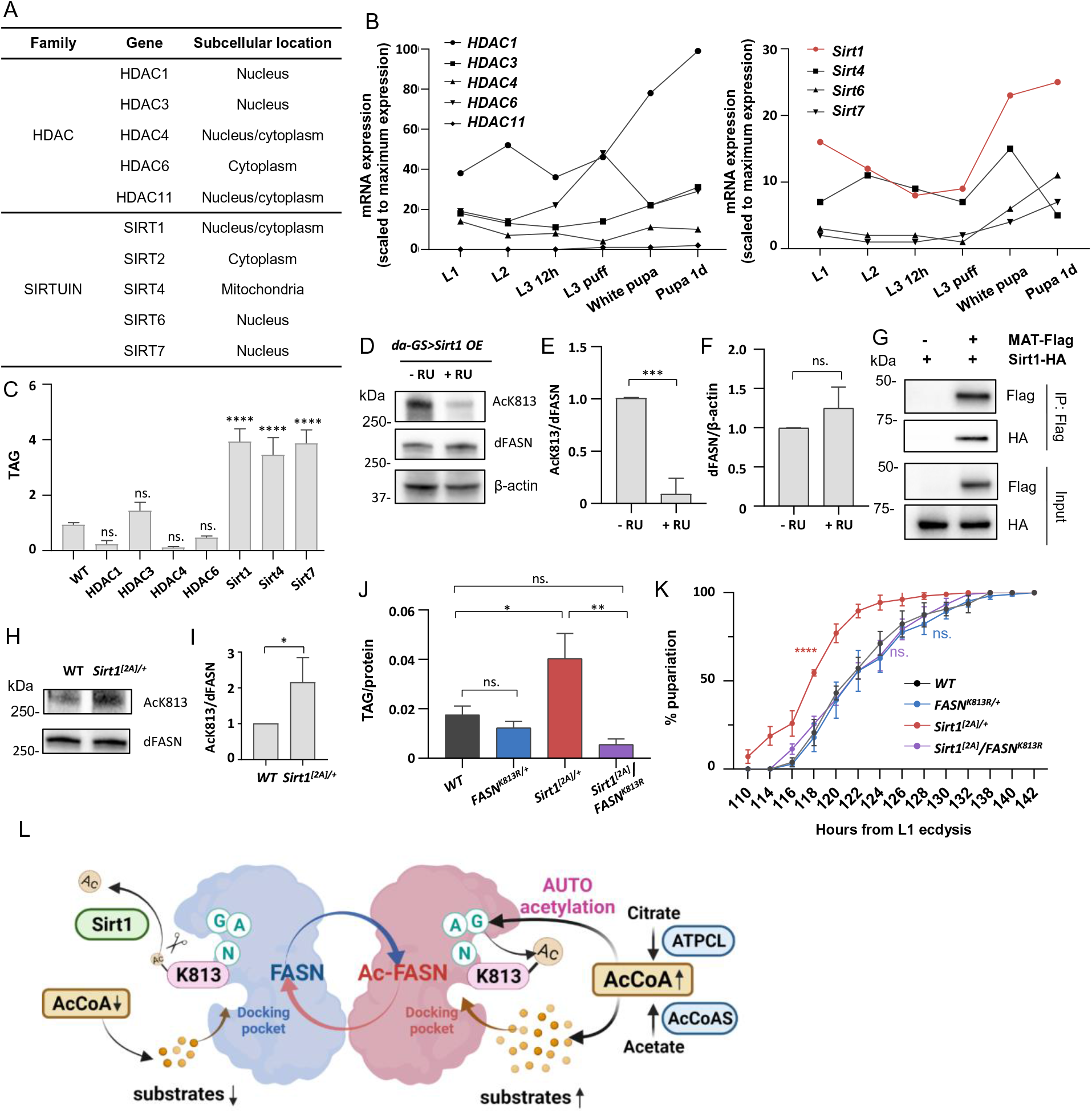
Sirt1-mediated deacetylation of dFASN1 regulates lipogenesis and developmental timing in *Drosophila* larvae. (A) Table summarizing known *Drosophila* KDACs and their subcellular localizations. (B) Developmental mRNA expression of KDACs. The transcriptional profiles of KDACs were retrieved from the *Drosophila* developmental transcriptome study (Graveley, Brooks et al. 2011). (C) TAG levels of KDAC mutants at L3 stage. TAG was normalized against total protein amount. Y-axis represents folds change *vs.* WT (*yw^R^ or w^1118^*). One-way ANOVA (*vs. WT*). Error bars represent SD; \*\*\*\**p*<0.0001; \*\**p*<0.01; \**p*<0.05; ns. *p*>0.05. (D) Acetylation of K813 in L3 larvae overexpressing *Sirt1.* Western blotting was performed using anti-AcK813. Daughterless GeneSwitch (GS)-Gal4 driver was used to drive ubiquitous expression of *Sirt1*. The expression was activated by 100 µM of RU468 (+ RU). (E) Quantification of K813 acetylation in panel D. K813 acetylation was normalized against total dFASN. Error bars represent SD. *t test*. \*\*\**p*<0.001. (F) Quantification of dFASN expression in panel D. dFASN expression was normalized against β-actin. Error bars represent SD. *t test*. ns. *p*>0.05. (G) Co-immunoprecipitation of Sirt1 and the MAT domain of dFASN. Kc167 cells were transfected by plasmids encoding Flag-tagged MAT and HA-tagged Sirt1. Interaction between MAT and Sirt1 was determined by western blotting analyses with anti-HA antibodies. (H) Acetylation of K813 in *WT* (*yw^R^/w^1118^*) and *Sirt1^[2A]/+^* larvae. (I) Quantification of K813 acetylation in panel H. K813 acetylation was normalized against dFASN. Error bars represent SD. *t test*. \**p*<0.05. (J) TAG levels in *WT*, *FASN^K813R^*^/+^, *Sirt1^[2A]/+^*, and *Sirt1^[2A]^*/*FASN^K813R^.* Error bars represent SD. One-way ANOVA (*vs*. *WT*). \**p<*0.05; \*\**p<*0.01; ns. *p*>0.05. (K) Developmental timing of *yw^R^/w^1118^* (*WT*), *FASN^K813R^*^/+^, *Sirt1^[2A]/+^*, and *Sirt1^[2A]^*/*FASN^K813R^.* Number of pupariation was counted every 2-4 hours. Log-rank test (*vs. WT*). Error bars represent SD; \*\*\*\**p*<0.0001; ns. *p*>0.05. (L) Working model showing acetyl-CoA-mediated autoacetylation of fatty acid synthase. The model figure was created with BioRender.com.

To determine whether Sirt1 regulates dFASN acetylation at K813, we ectopically expressed *Sirt1* using daughterless GeneSwitch-Gal4 driver (da-GS-Gal4) at the L3 stage. Remarkably, short-term overexpression (1 day) of *Sirt1* removed acetylation at K813 by about 90% **(Figure 7D-7E)**. Overexpression of *Sirt1* did not alter dFASN protein expression **(Figure 7D, 7F),** although it was reported that the transcription of FASN is indirectly regulated by *Sirt1* through deacetylation of SREBP1c (Ponugoti et al., 2010; Wang et al., 2010b; Xu et al., 2010b). We further showed that Sirt1 interacted with MAT domain of dFASN through a co-immunoprecipitation experiment by co-expressing HA-tagged Sirt1 and Flag-tagged MAT domain in *Drosophila* Kc167 cells (**Figure 7G**).

Sirt1 is a known negative regulator in lipid synthesis pathways (Walker et al., 2010; Xu et al., 2010a). Indeed, we found that body fat accumulation was increased in *Sirt1^[2A]/+^* loss-of-function mutants at L3 stage **(Figure 7J)**, accompanied by elevated acetylation of K813 **(Figure 7H-7I)**. To determine if K813 acetylation is required for Sirt1-mediated lipogenesis, we performed an epistasis analysis by measuring TAG levels of *Sirt1^[2A]^/FASN^K813R^* double mutants **(Figure 7J)**. Intriguingly, the elevated TAG levels in *Sirt1^[2A]/+^* was rescued by *FASN^K813R/+^* mutants, suggesting that the role of Sirt1 in body fat accumulation is through dFASN deacetylation. Consistently, *Sirt1^[2A]/+^* developed faster and showed early pupariation compared to wild-type flies, which was rescued by *FASN^K813R/+^* mutants **(Figure 7K)**. Taken together, these results demonstrate that Sirt1 is the primary deacetylase targeting K813. Acetylation of K813 is required for Sirt1-mediated lipogenesis and larval development.

## DISCUSSION

Metabolic homeostasis plays an important role in animal development and growth (Hochachka, 2003; Leopold and Perrimon, 2007). One novel mechanism underlying the coordination of metabolic homeostasis and growth is the interplay between metabolic intermediates and protein post-translational modifications (Drazic et al., 2016; Lu and Thompson, 2012). Among all metabolic intermediates, acetyl-CoA has emerged as a central player that links metabolism to cellular signaling, especially via lysine acetylation of histone and non-histone proteins (Pietrocola et al., 2015; Shi and Tu, 2015). In the present study, we uncover a novel role of acetyl-CoA-mediated autoacetylation of dFASN in lipogenesis during *Drosophila* larval development (**Figure 7L**). On the one hand, acetyl-CoA fuels dFASN as the carbon donor for the growing fatty acid chain. On the other hand, acetyl-CoA, as the acetyl-group donor, directly modulates dFASN enzymatic activity through acetylation of the critical lysine residue K813. Additionally, we surprisingly found that acetylation of dFASN does not require KATs; instead, it is mediated by a conserved P-loop-like motif ‘N-xx-G-x-A’ neighboring K813 (**Figure 7L**). Lastly, we identified Sirt1 as the primary deacetylase for dFASN, which acts as a negative regulatory mechanism (**Figure 7L**). In summary, we discovered a previously unknown regulatory mechanism for metabolic homeostasis in developing animals that links acetyl-CoA, autoacetylation, and DNL.

### Acetylation of dFASN at K813 as a novel fine-tune mechanism for developmental DNL and metabolic homeostasis

Under conditions like excess nutrition, growth factor stimulation, obesity, or cancer, DNL is significantly elevated, which is thought to be mainly controlled through the transcriptional activation of lipogenic enzymes (Strable and Ntambi, 2010). The mRNA expression of FASN is positively correlated with elevated lipogenesis in conditions like obesity, diabetes, fatty liver diseases, and cancers (Angeles and Hudkins, 2016; Kridel et al., 2007; Loftus, 2000; Wu et al., 2011). However, the protein levels of FASN are rarely characterized in these studies. Interestingly, several conflicting results show little correlation between FASN protein expression and its enzymatic activity (Hennigar et al., 1998; Jin et al., 2010; Najjar et al., 2005; Qureshi et al., 1975; Sabbisetti et al., 2009). Consistent with these studies, we found that the protein levels of dFASN remain unchanged during *Drosophila* larval development, which does not correlate with dFASN enzymatic activity. In contrast, we find that acetylation modification of dFASN at lysine K813 is positively associated with dFASN activity and developmental lipogenesis. Our findings suggest that lysine acetylation of dFASN, but not translational regulation, plays a crucial role in dFASN-mediated lipogenesis.

Indeed, our genetic analysis further demonstrates that K813R substitution reduces dFASN enzymatic activity and lipogenesis, while acetyl-CoA-mediated acetylation of recombinant dFASN proteins increases enzyme activity. Although acetylation of FASN has been reported in several previous global acetylome studies, the functional roles of FASN acetylation remain largely unknown. A recent study investigated the role of FASN acetylation in DNL in human cell culture (Lin *et al*., 2016). The study shows that treatment of KDAC inhibitors induces the acetylation of hFASN, promotes FASN degradation, and reduces lipogenesis. However, it remains to be determined whether the regulation of lipogenesis by KDAC inhibition is due to global acetylation, or it is directly through FASN acetylation. In addition, the functional lysine residues of hFASN that are responsible for altered lipogenesis are not identified. Because of the high conservation between K813 of dFASN and K673 of hFASN, it is possible that K673 is the key lysine residue mediating DNL of human cells.

Apart from K813, three other lysine residues of dFASN (K926, K1800, and K2466) are highly conserved among animal species, and their homologs are also found to be acetylated in other animal species. Because these lysine residues are located on different domains, it is not hard to imagine that acetylation of each lysine may play distinct roles related to their associated domains. In the present study, we show that acetylation of K813, but not K926, modulates dFASN activity, body fat accumulation, and *Drosophila* developmental timing. It is likely that acetylation of K926 affects other aspects of enzyme properties and functions that are less important for larval development. Different from K926, K813 is at the substrate docking pocket of the MAT domain. This unique localization suggests that acetylation of K813 might introduce conformational changes of the docking site and modulate dFASN catalytic activity in response to substrate availability during larval development.

### **N-xx-G-x-A** motif as a novel mechanism for rapid autoacetylation

Another surprising finding from our study is that acetylation of dFASN at K813 does not require a known KAT; rather it is autoacetylated by acetyl-CoA in a dosage-dependent manner. Acetyl-CoA is an important metabolic intermediate that plays dual roles in maintaining metabolic homeostasis, as metabolic precursors in energy metabolism and acetyl-group donor for protein acetylation. The cytosolic pools of acetyl-CoA increase under feeding or excess nutrient conditions (Shi and Tu, 2015). Consistently, our studies reveal that the amount of intracellular acetyl-CoA elevates in fast-growing larvae, which could modulate dFASN activity by promoting both the biosynthesis of malonyl-CoA and autoacetylation of lysine residue K813 for the conformational changes of malonyl-CoA docking pocket.

It was previously thought that only mitochondria proteins were nonenzymatically acetylated since no KATs have been identified in mitochondria. Besides, the high acetyl-CoA concentration and relatively high pH of the mitochondrial matrix facilitate the lysine nucleophilic attack on the carbonyl carbon of acetyl-CoA (Wagner and Payne, 2013). While on contrast, the cytosolic acetyl-CoA level is much lower, and the pH is neutral. Therefore, it has been postulated that acetylation of cytosolic protein is catalyzed by various cytoplasm-localized KATs. Recently, KAT-independent acetylation of cytosolic proteins has been reported (Kulkarni et al., 2017; Olia et al., 2015). Yet, the underlying mechanism for nonenzymatic acetylation, especially of cytosolic proteins, remains largely unknown. When investigating how dFASN is autoacetylated by acetyl-CoA, we uncover a novel motif N-xx-G-x-A near acetylated K813. Substituting any of the three key amino acids largely blocks acetyl-CoA-mediated dFASN autoacetylation. The N-xx-G-x-A motif resembles the signature P-loop sequence (Q/R-xx-G-x-A/G) of KATs, which is required for acetyl-CoA recognition and binding (Roth et al., 2001). We predict that the N-xx-G-x-A motif of dFASN performs a similar function as the P-loop of KATs for acetyl-CoA binding, the key step in dFASN autoacetylation. Moreover, the N-xx-G-x-A motif is highly conserved among almost all FASN proteins, pointing out a conserved mechanism for autoacetylation of FASN.

In addition to the well-established KATs of the MYST, p300/CBP, and GCN5 families, there are over 15 proteins that have been reported to possess KATs activity (Sheikh and Akhtar, 2019), such as CLOCK (Doi et al., 2006) and Eco1 (Ivanov et al., 2002). Since FASN may contain an acetyl-CoA binding motif of KATs, it is possible that FASN, particularly MAT domain, possesses KATs activity and may acetylate other proteins, especially those in DNL pathways. This possibility may be further explored through acetylome analysis in the future.

In summary, we uncover a previously unappreciated role of FASN acetylation in developmental lipogenesis and a novel mechanism for lysine autoacetylation. Our findings provide new insights into acetyl-CoA-mediated metabolic homeostasis during animal development. In addition, our studies underscore a promising therapeutic strategy to combat metabolic disorders by targeting autoacetylation of FASN.

## AUTHOR CONTRIBUTIONS

Conceptualization, T. M., and H. B.; Methodology, T. M., and H. B.; Validation, T. M.; Formal Analysis, T. M.; Investigation, T. M., J. K., P. K., and H. B.; Resources, H. B.; Writing – Original Draft, T. M; Writing –Review & Editing, H. B., and T. M.; Visualization, T. M., and H. B.; Supervision, H. B.; Project Administration, H. B., and T. M.; Funding Acquisition, H. B., and T. M.

## ACKNOWLEDGMENTS

We thank Bloomington *Drosophila* Stock Center and Drosophila Genomics Resource Center for fly stocks and cDNA clones. We thank BestGene Inc for *Drosophila* embryo injection service. We thank Ross Tomaino from Harvard Medical School Taplin Mass Spectrometry Facility for mass spectrometry analysis. We thank Ann Perera and Lucas Showman from ISU W. M. Keck Metabolomics Research Laboratory for mass spectrometry and LC-MS/MS analysis. We thank Baoyu (Stone) Chen and Sheng Yang for the DNA clones and reagents and the help with recombinant protein expression. We thank Justin Walley for the mass spectrometry analysis. We thank Basil Nikolau for the help with data interpretation. Graphical abstract and working model figures were created with BioRender.com. The protein structures were visualized with PyMOL Molecular Graphics System. This work was supported by NIH R01AG058741 to H.B., Glenn/AFAR Scholarships to T.M.

## DECLARATION OF INTERESTS

The authors declare no competing interests.

## Supplemental information

**Figure S1.**
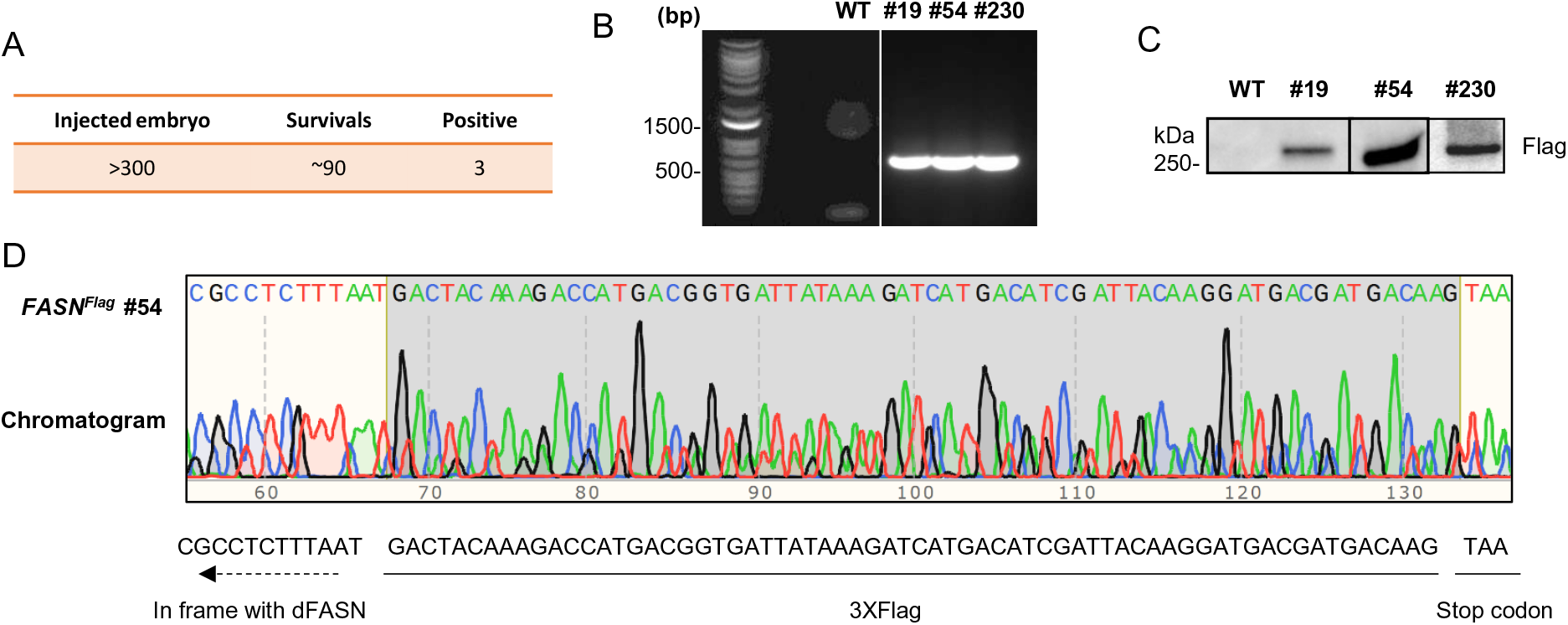
Generation of Flag knock-in flies. (A) Summary table for the number of injected fly embryos, survived injections, and successful Flag knock-in lines. (B) Positive knock-in fly lines were verified by PCR with a forward primer (5’-3’: GCAGCTTCAAGCAGCGTTAC) recognizing *dFASN* sequence and a reverse primer (5’-3’: CACCGTCATGGTCTTTGTAGTC) recognizing Flag sequence. (C) Positive knock-in fly lines verified by western blot with anti-Flag. (D) Positive knock-in line #54 was verified by Sanger sequencing. This line was used for endogenous dFASN pull-down in this study.

**Figure S2.**
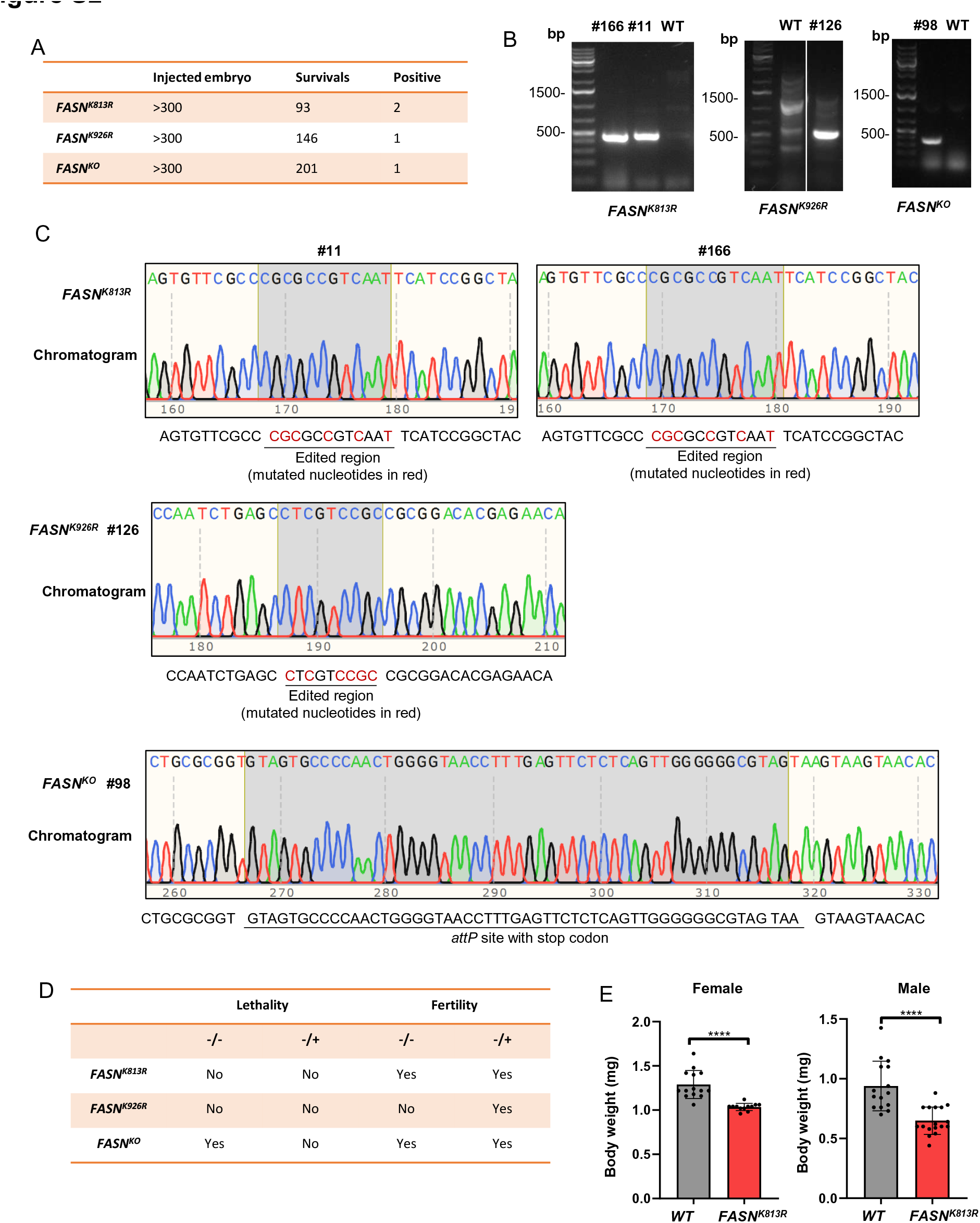
Generation of dFASN acetylation deficient and knockout mutants. (A) Summary of the isolation of dFASN acetylation deficient fly mutants (*FASN^K813R^* and *FASN^K926R^*) and knockout fly mutant *FASN^KO^*. The number of injected embryos, survived injections, and positive mutant lines are summarized in the table. (B) PCR verification of *FASN* mutants. *FASN^K813R^* mutants were verified by PCR with a forward primer (5’-3’: ttcgccCGCgcCgtCaaTtc, mutated nucleotides are in lowercases) recognizing *FASN^K813R^* mutated sequence and a reverse primer (5’-3’: AGCTGATAGGACGCACTAGA) recognizing *dFASN* sequence. The correct PCR product size was 448 bp. *FASN^K926R^* mutants were verified by PCR with a forward primer (5’-3’:AATCTGAGCcTcGTccgcCG, mutated nucleotides are in lowercases) recognizing *FASN^K926R^* mutated sequence and a reverse primer (5’-3’: GCCAAGCTACCGCTCTCACA) recognizing *dFASN* sequence. The correct PCR product size was 502 bp. *FASN^KO^* mutants were verified by PCR with a forward primer (5’-3’:CCTTTGAGTTCTCTCAGTTG) recognizing *attP* sequence and a reverse primer (5’-3’: CGTATCCATTGCCAGACTCAT) recognizing *dFASN* sequence. The correct PCR product size was 374 bp. (C) Successfully mutated fly lines verified by Sanger sequencing. (D) Fertility and lethality of dFASN acetylation deficient fly mutants (*FASN^K813R^* and *FASN^K926R^*) and knockout fly mutant *FASN^KO^*. (E). Body weight of *WT* (*yw^R^*) and *FASN^K813R^*, 3-day old adult flies. *t*-test. \*\*\*\**p*<0.0001.

**Figure S3.**
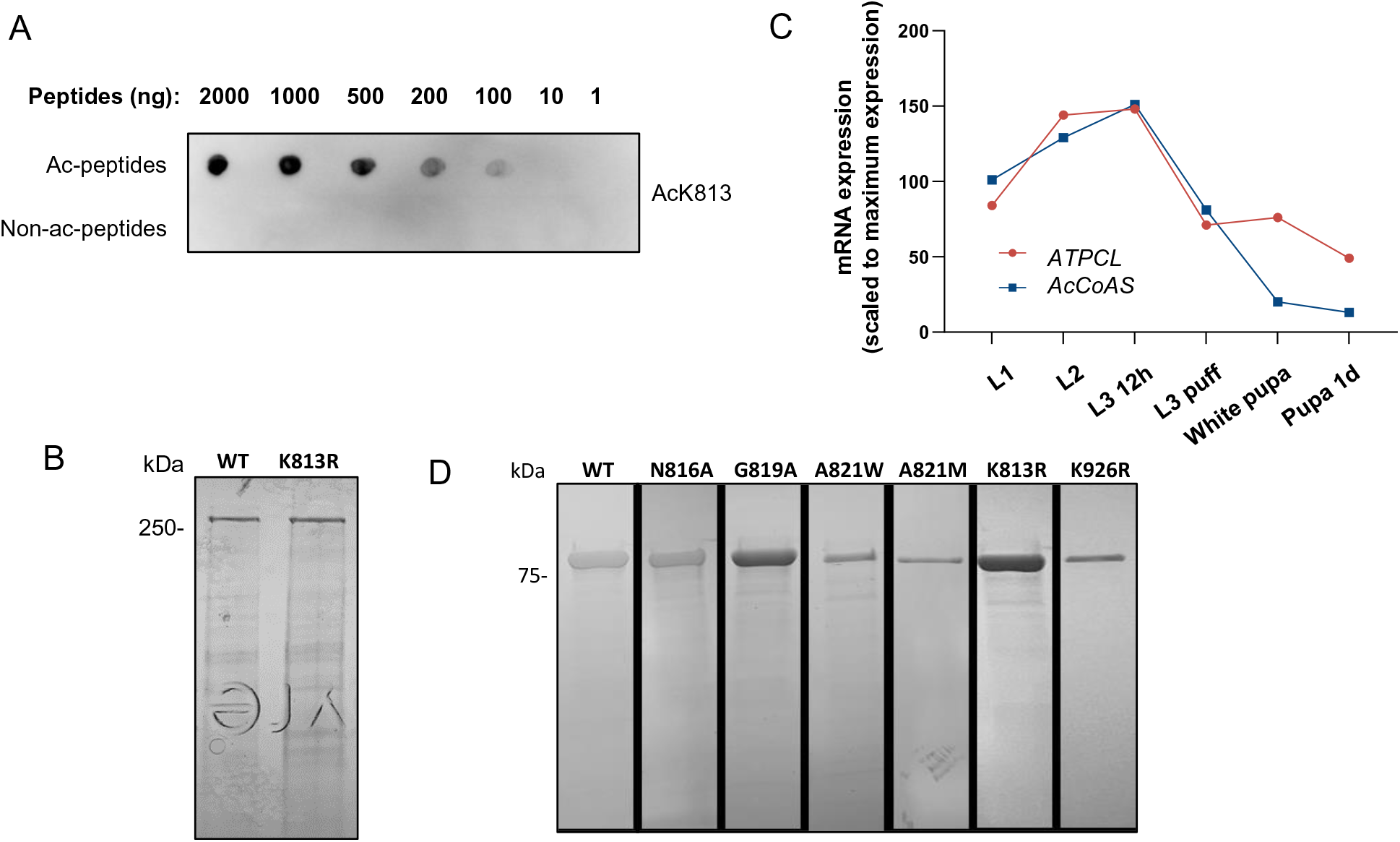
Expression of recombinant dFASN and the expression pattern of *ATPCL* **and *AcCoAS***. (A) Specificity of the acetylated antibody (anti-AcK813) determination by dot-plot. Ac-peptides: acetylated lyophilized peptides (NAEGVFA-acK-AVNSSG); Non-ac-peptides: non-acetylated lyophilized peptides (NAEGVFAKAVNSSG). (B) Coomassie Blue staining of purified WT and K813R recombinant dFASN protein from *Sf9* cells. Purified recombinant dFASN proteins were around 270 kDa. (C) Developmental mRNA expression of *ATPCL* and *AcCoAS*. The transcriptional profiles of KATs were retrieved from the *Drosophila* developmental transcriptome study (Graveley, Brooks et al. 2011). (D) Coomassie Blue staining of WT and mutants recombinant proteins from *E.coli* BL21 (DE3) cells. Purified recombinant KS-MAT proteins were around 90 kDa.

## STAR*METHODS

### KEY RESOURCES TABLE

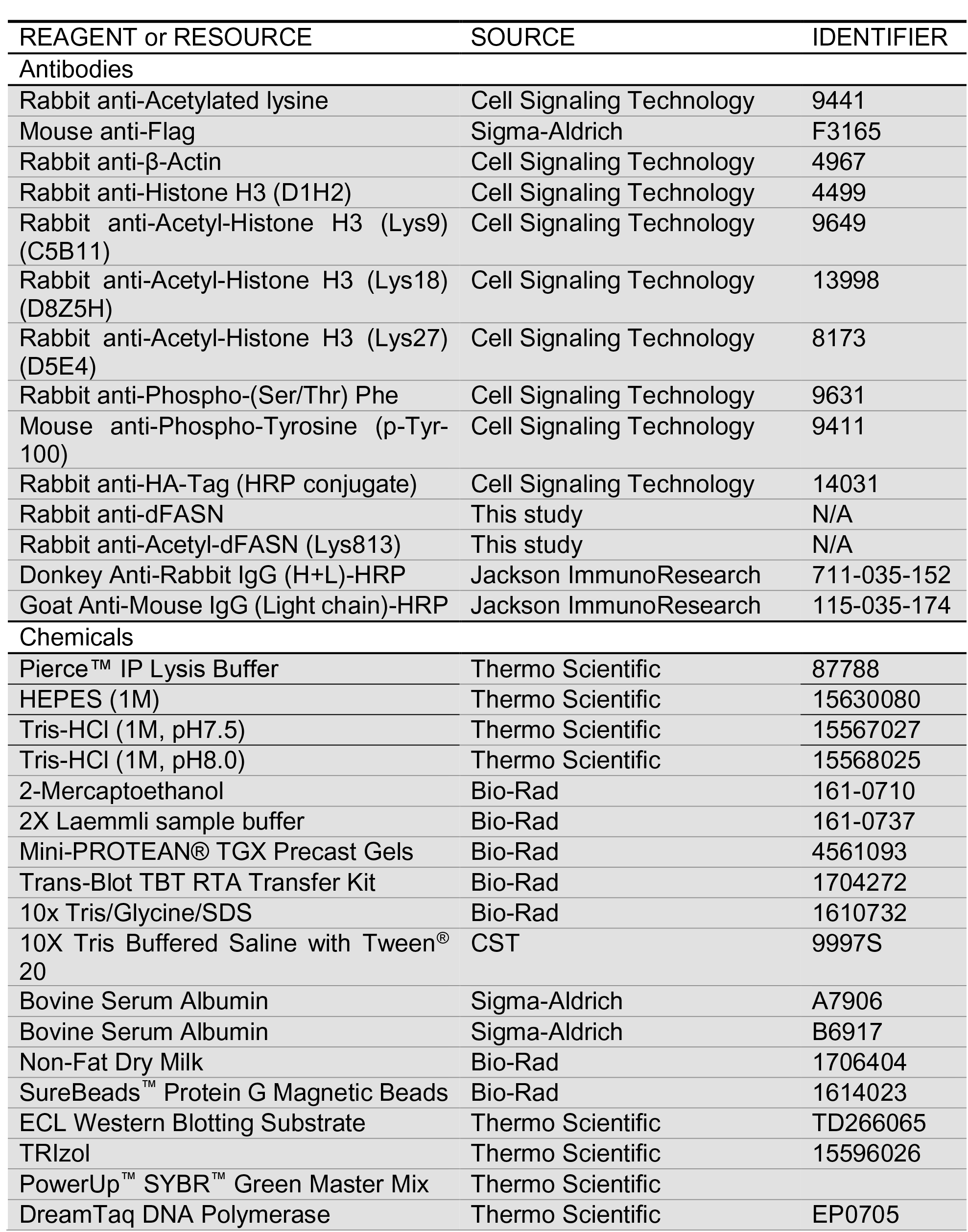
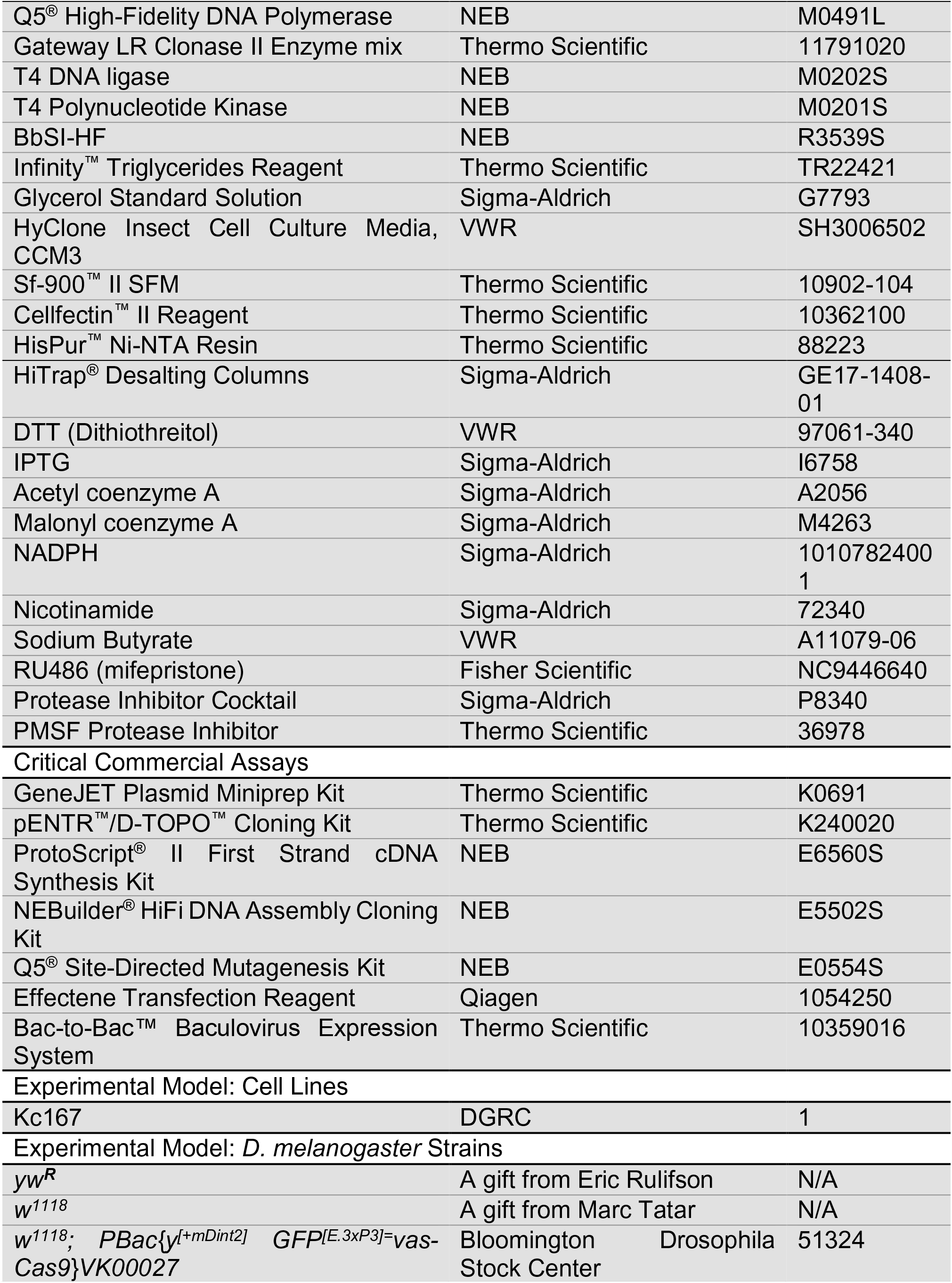
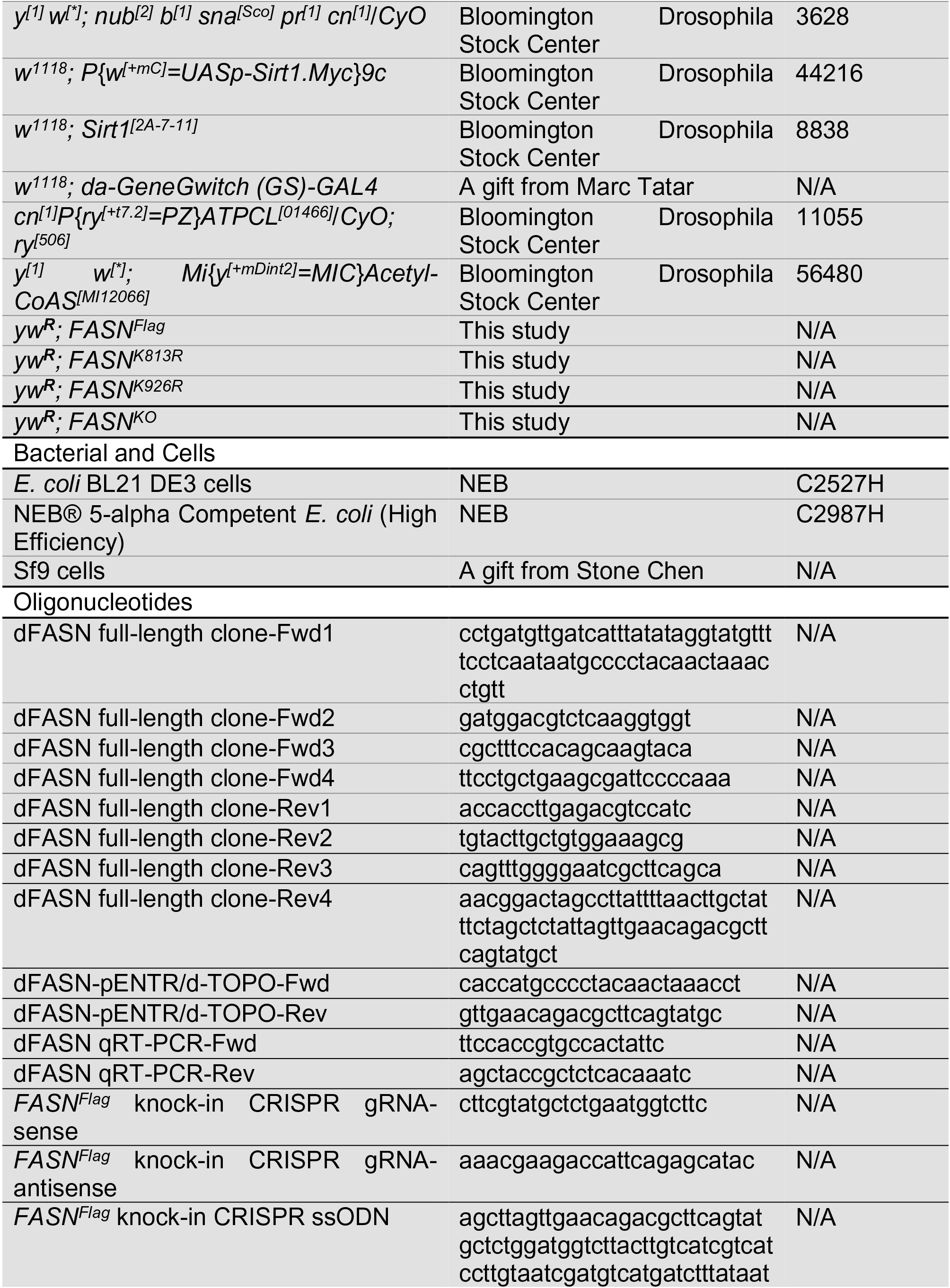
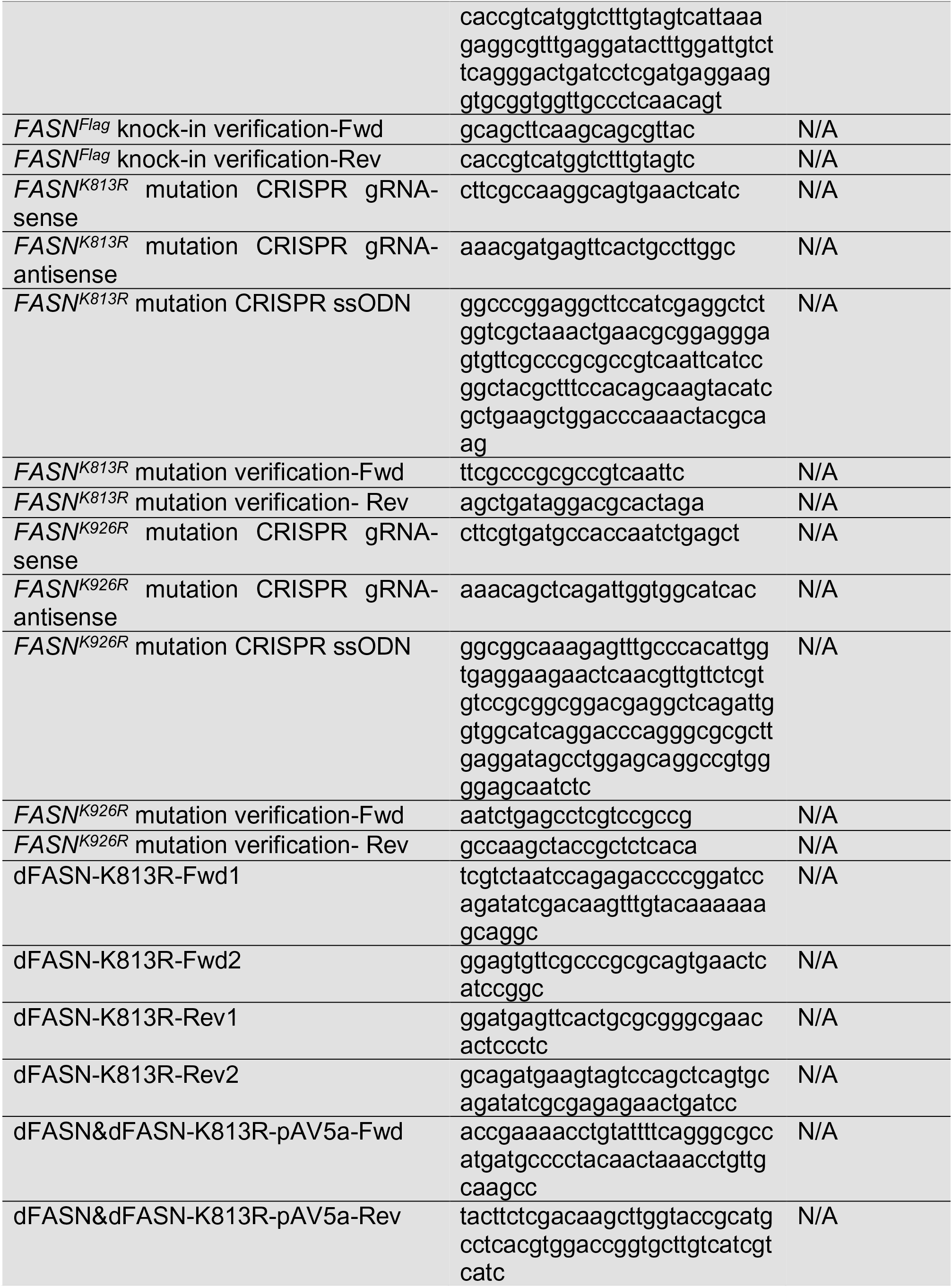
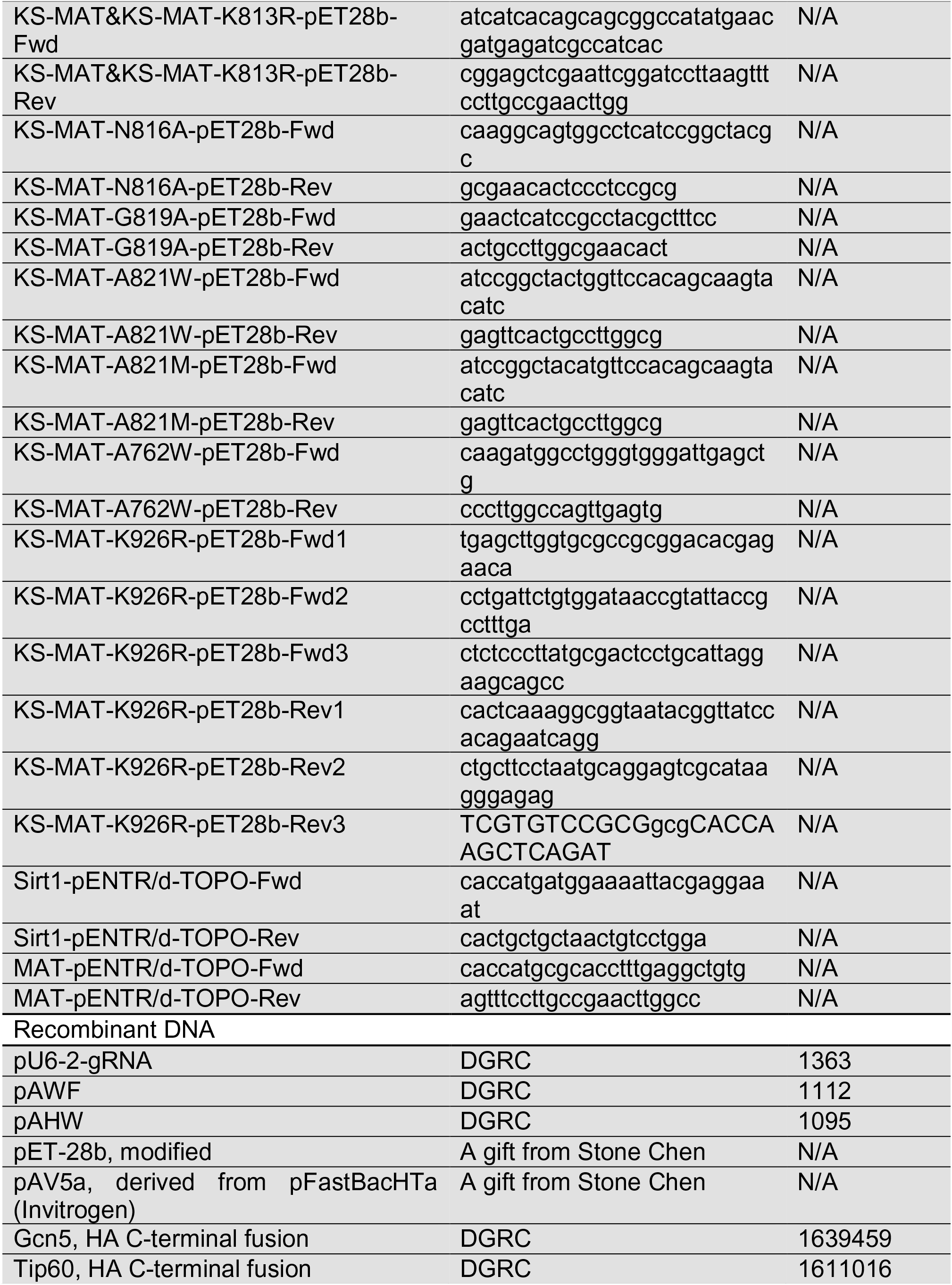
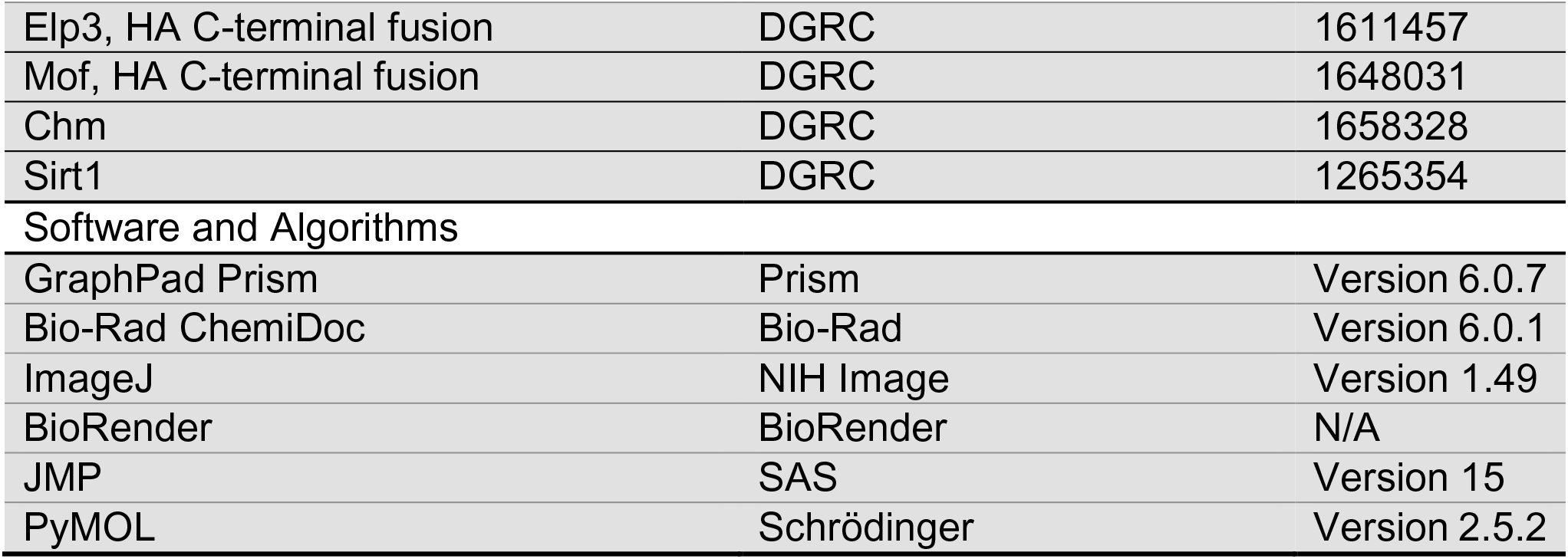

## RESOURCE AVAILABILITY

### Lead contact

Further information and requests for reagents and resources should be directed to and will be fulfilled by the Lead Contact Hua Bai.

### Materials availability

Plasmids and fly knock-in or mutant lines generated in this study are available from Dr. Hua Bai.

### Data and code availability

Mass spectrometry datasets generated in this study are available upon request.

## EXPERIMENTAL MODEL AND SUBJECT DETAILS

### Cell Culture and Plasmid Construction

*D. melanogaster* Kc167 cells were grown in CCM3 cell culture medium in an incubator at 25°C. cDNA clones for Gcn5, Tip60, Elp3, and Mof (with HA tag) were purchased from DGRC. cDNA clones for Chm and Sirt1 were purchased from DGRC and cloned into pAHW vector through pENTR/d-TOPO and Gateway LR Ligation. Full-length cDNA of dFASN, KS-MAT didomain, and MAT domain was cloned from *w^1118^* flies by HIFI Assembly. The gene coding for the MAT domain was cloned into pAWF vector through pENTR/d-TOPO and Gateway LR Ligation. For recombinant protein expression, full-length dFASN and KS-MAT didomain were cloned into pAV5a (derived from pFastBacHTa) and pET28b vector, respectively. Single amino acid mutations were introduced to those plasmids by HIFI Assembly or Q5^®^ Site-Directed Mutagenesis kit.

### *Drosophila* Husbandry and Strains

A detailed list of fly strains is provided in the Key Resources Table. *yw^R^* was used as the control in the TAG measurement, developmental timing, and FASN activity assay, unless otherwise noted in the figure legend.

Flies were maintained at 25 ^0^C, 60% relative humidity, and a 12-hour light/dark cycle. Adults and larvae were reared on standard cornmeal and yeast-based diet. The standard cornmeal diet consists of the following materials: 0.8% cornmeal, 10% sugar, and 2.5% yeast. For developmental timing analysis, adult flies were maintained in small cages and allowed to lay eggs for 4-6 hours on apple juice agar plates supplemented with yeast paste. Newly hatched larvae were transferred to vials with standard *Drosophila* medium. For the use of inducible Gal4/UAS system, the induction of transgene expression was achieved by feeding larvae on RU486 food for one day. RU486 was dissolved in 95% ethanol and added to standard food at a final concentration of 100 µM.

## METHOD DETAILS

### Precise Genome-editing

Flag knock-in fly line (*yw; FASN^Flag^*) was generated by CRISPR/Cas9-mediated homology-directed repair by ssODNs (Boel et al., 2018). Briefly, the gRNA with a PAM site within 10 nucleotides from the site of editing was designed according to the Fly CRISPR Optimal Target Finder (Gratz et al., 2014). The sense and antisense gRNA oligos were annealed and then ligated to the pU6-2-chiRNA vector that expressed chimeric RNA under the control of the *Drosophila* U6 promoter. The ssODN with two homology arms and the edited region (the last 34 nucleotides of dFASN with silent mutations and 3XFlag sequences) was ordered from Integrated DNA Technologies. The pU6-2-chiRNA and ssODN were injected into the embryos of vas-Cas9 flies (*w^1118^; PBac{y^[+mDint2]^ GFP^[E.3xP3]=^vas-Cas9}VK00027*) by BestGene Inc. dFASN acetylation-deficient fly lines (*yw; FASN^K813R^* and *yw; FASN^K926R^*) and knockout fly line (*yw; FASN^KO^*) were generated by the same approaches. dFASN knockout fly line was generated by inserting an *attP* sequence and premature stop codon to 2^nd^ exon of dFASN.

### Antibody Generation

The rabbit polyclonal antibodies against total dFASN and acetylated K813 were generated by immunizing rabbits with synthetic peptides, followed by affinity purification (Yenzym antibodies, CA, USA). The antibody for dFASN was raised against the peptide NAEGVFAKAVNSSG, which covers amino acids 806-819 of the dFASN protein. The antibody against acK813 was raised against the peptide NAEGVFA-acK-AVNSSG.

### Developmental Timing

Flies were allowed to lay eggs on apple juice agar petri dishes with yeast paste for 3 hours. The plates were then maintained in a fly incubator for 20-24 hours. To synchronize the animals, newly hatched L1 larvae were transferred to fly vials with standard food (15-30 larvae/vial). Larvae were maintained in the fly incubator, and the number of pupae was counted every 2-4 hours.

### Immunoprecipitation

Fly larvae or Kc167 cells were lysed with Pierce^™^ IP Lysis Buffer supplied with deacetylase inhibitor nicotinamide at 4℃ for 20 min. After the removal of unbroken cells and debris by centrifugation (14,000 rpm/30 min), the soluble fractions were collected and incubated with mouse anti-Flag M2 antibody (1:100) at 4℃ overnight. The next day, the lysate-antigen mixture was incubated with SureBeads^™^ Protein G Magnetic Beads at 4℃ for 3 hours. The magnetic beads were then washed three times with Pierce^™^ IP Lysis Buffer. Proteins were denatured and eluted from the beads with Laemmli sample buffer at 95 ℃ for 5 min.

### Western Blot Analysis

Fly larvae were lysed with 20 mM Tris-HCl (pH 8.0), 100 mM NaCl buffer supplied with deacetylase inhibitor nicotinamide at 4℃ for 20 min. After centrifugation at 14,000 rpm for 30 min, the soluble fractions were collected and denatured with Laemmli sample buffer at 95℃ for 5 min. Proteins were separated by Mini-PROTEAN^®^ TGX Precast Gels, transferred to PVDF membranes, immunoblotted with the indicated antibodies, and visualized with Pierce ECL Western Blotting Substrate. For detailed antibody information, see Key Resources Table.

### Acetylation Site Identification by Mass Spectrometry

Larvae of dFASN-Flag knock-in flies were lysed, and endogenous dFASN-Flag was precipitated as described above. Proteins were run and separated on an SDS-PAGE gel, followed by Coomassie Blue staining. The protein band of dFASN at around 270 kDa was excised and the acetylation sites were identified by mass spectrometry analysis at the Taplin Biological Mass Spectrometry Facility of Harvard Medical School.

### Quantification of Acetyl-CoA by LC-MS/MS

Acetyl-CoA levels were measured by LC-MS/MS as previously described (Hu et al., 2003). About 300 mg of L2 and L3 larvae were snap-frozen in liquid nitrogen and pulverized into powder with mortar and pestle. 200 μL/100 mg of solvent buffer (5% sulfosalicylic acid containing 50 μM DDT) was added to the sample immediately. Samples were sonicated three times for 10 s on ice, and the supernatant was collected by centrifugation at 14,000 xg for 20 min. Right before LC-MS/MS analysis, 2 μL of Ammonia (25%) was added to 98 μL of the sample solutions. LC-MS/MS analysis was performed at W. M. Keck Metabolomics Research Laboratory of Iowa State University following the method described previously (Hu et al., 2003).

### Quantification of TAG

TAG quantitation was performed with Infinity^™^ Triglycerides Liquid Stable Reagent. Briefly, 20-30 mg of fly larvae were collected and homogenized in 150 µL 1X PBS containing 0.05 % Tween 20. A particle-free supernatant solution was obtained by centrifugation at 14,000 xg for 15 min. 10 μL of the homogenized sample was removed for protein content measurement with a Bradford assay. The rest of the sample was heated for 10 min at 70℃. 20 μL of supernatant or glycerol standards was added to 200 µL of Triglycerides Liquid Stable Reagent. The absorbance at 550 nm was read with the plate reader at 37℃. TAG content of each sample was normalized against protein concentrations.

### FASN Activity Assay

FASN activity assay was modified from previous studies (Dils and Carey, 1975; Lin et al., 2016; Menendez et al., 2004). In brief, fly larvae were homogenized with Dounce homogenizer in lysis buffer, 20 mM HEPES (pH 7.2), 250 mM sucrose, 2 mM MgCl_2_, 1mM dithiothreitol (DTT), and 1mM EDTA, supplied with protease inhibitor cocktail. A particle-free supernatant solution was obtained by centrifugation at 14,000 xg for 15 min. Protein concentration was determined by Pierce^™^ Coomassie Plus (Bradford) Assay. 20 µL of protein extract was added to 200 µL assay buffer containing 200 mM potassium phosphate buffer (pH 6.6), 1 mM DTT, 1 mM EDTA, 240 µM NADPH, and 31 µM acetyl-CoA. The reaction was monitored at 340nm, 25℃ for 2 min with the plate reader to measure background NADPH oxidation. After adding 90 µM of malonyl-CoA, the reaction was assayed for an additional 10 min to determine dFASN-dependent oxidation of NADPH. Oxidation of NADPH in each sample was calculated based on the NADPH-equivalent standard curve. FASN activity was calculated in nmol NADPH oxidized min^-1^ mg protein^-1^.

### Recombinant Protein Expression and Purification

The two constructions, dFASN-pAV5a and dFASN-K813R-pAV5a, were utilized to produce dFASN recombinant protein in the Bac-to-Bac protein expression system. The expression and purification methods of recombinant dFASN proteins were adapted from a previous study (Carlisle-Moore et al., 2005). Briefly, Sf9 cells were transfected with bacmid DNA. The virus was harvested from cell culture at 96 hours post-transfection. Both viral stocks were subjected to two rounds of amplification, after which the viral titer was determined. 800 mL of Sf9 cells were infected with recombinant baculovirus, and a cell pellet was collected 80 hours post-infection. The pellet was resuspended in 50 mL of lysis buffer containing 20 mM Tris-HCl (pH 8), 500 mM NaCl, 5 mM imidazole, and 20% glycerol, and frozen in -80℃ for overnight. The cell lysate was thawed on iced water and then centrifuged at 20, 000 xg for 50 min. The supernatant was applied to HisPur^™^ Ni-NTA Resin (5-ml bed volume) equilibrated with lysis buffer. The His-bind resin column was washed with wash buffer containing 20 mM Tris-HCl (pH 8), 500 mM NaCl, 60 mM imidazole, and 20% glycerol. Protein elution was performed using 20 mM Tris-HCl (pH 8), 200 mM NaCl, 500 mM imidazole, and 20% glycerol. Fractions containing wild-type or K813R proteins were then immediately exchanged to 0.8 M potassium phosphate buffer (PPB) (pH 7.4) containing 1 mM EDTA and 10 mM DTT. Protein expression and purification were analyzed by SDS-PAGE and Coomassie Blue staining. Protein concentration was determined by the absorbance at 280 nm using an extinction coefficient of 228390 M^-1^ cm^-1^ calculated at Expasy (www.expasy.ch). 100 µL aliquots of protein were snap-frozen in liquid nitrogen with 20% glycerol for storage at -80℃. Protein yield was assessed using the calculated molecular mass of 278 kDa (www.expasy.ch).

KS-MAT recombinant proteins were expressed in *E. coli* BL21 (DE3) cells (Rittner et al., 2020; Rittner et al., 2018). Wild-type or mutant constructs were transformed into chemically competent cells. The transformants were grown for 16 hours at 37℃ in 20 mL of start culture in LB medium supplied with 100 µg/mL kanamycin and 1% glucose. Pre-cultures were used to inoculate 500 mL LB medium (100 µg/mL kanamycin and 0.2% glucose). Cultures were grown at 37℃ until they reached an optical density (OD600) of 0.5–0.6. After cooling at 4℃ for 20 min, cultures were induced with 0.25 mM IPTG, and grown for additional 16 hours at 20℃ and 180 rpm. Cells were harvested by centrifugation (3,900 rpm for 30 min). Cell pellets were resuspended by 10 mL of lysis buffer containing 50 mM PPB (pH7.4), 200 mM KCl, 20 mM imidazole, 1 mM EDTA, 1 mM DTT, and 10% glycerol, and were sonicated at 20% amplification for 8 x 15s bursts with 30s off between bursts. Lysates were cleared by centrifugation at 20, 000 xg for 30 min and were mixed with 1 M MgCl_2_ to a final concentration of 2 mM. Similar His-binding steps were performed as described above, except that the resin columns were washed with wash buffer containing 50 mM PPB (pH7.4), 200 mM KCl, 20 mM imidazole, 1 mM DTT, and 10% glycerol. Proteins were eluted with elution buffer containing 50 mM PPB (pH7.4), 200 mM KCl, 400 mM imidazole, and 10% glycerol. Fractions containing wild-type or mutant proteins were immediately exchanged to storage buffer containing 250 mM PPB (pH 7.4), 1 mM EDTA, 1mM DTT, and 10% glycerol. Protein expression and purification were analyzed by SDS-PAGE and Coomassie Blue staining. Protein concentration was determined by the absorbance at 280 nm using extinction coefficients calculated at Expasy (www.expasy.ch). 100 µL aliquots of protein were snap-frozen in liquid nitrogen for storage at -80℃. Protein yield was assessed using the calculated molecular mass of (www.expasy.ch).

### *In vitro* Acetylation Assay

Acetyl-CoA was diluted to different concentrations with assay buffer containing 20 mM Tris-HCl (pH7.0 or 8.0), 500 mM NaCl. Recombinant proteins and acetyl-CoA were mixed in assay buffer to a final volume of 40 µL. The reactions were incubated in a water bath at 37℃ for 1 hour or 3 hours. BSA was dissolved in assay buffer (pH7.0 or pH8.0), and then incubated with acetyl-CoA at the same condition for 6 hours. The reactions were terminated by adding Laemmli sample buffer and heat-shock at 95℃ for 5 min. Acetylation was analyzed by western blotting with acetylated antibodies.

### Kinetics Evaluation

The assay condition described in FASN activity assay was also used for the determination of kinetics parameters. The enzymatic activity of WT dFASN or K813R mutants upon varying concentration of malonyl-CoA was determined at fixed concentrations of NADPH (240 µM) and acetyl-CoA (31 µM). For the kinetics evaluation with acetyl-CoA pre-incubation, acetyl-CoA was added to dFASN proteins to a final concentration of 20 µM. Proteins were then incubated at 37℃ for 1 hour, followed by dFASN activity assay. For control groups, dFASN proteins were mixed with 20 µM of acetyl-CoA right before the activity assay without any pre-incubation. Kinetics parameters were calculated by fitting the observed velocities to the Michaelis-Menten equation using GraphPad.

## QUANTIFICATION AND STATISTICAL ANALYSIS

GraphPad Prism and JMP were used for statistical analysis. Unpaired two-tailed Student’s *t-*test or ordinary one-way ANOVA was performed to compare the mean value between control and treatment groups. The outliers were excluded using robust regression and outlier removal method (Q = 10%) prior to the data analysis. Developmental timing data was fitted to Survival Model and analyzed by Log-rank test in JMP.

## Notes

### Competing Interest Statement

The authors have declared no competing interest.

